# KAT5 regulates neurodevelopmental states associated with G0-like populations in glioblastoma

**DOI:** 10.1101/2022.03.17.484768

**Authors:** Anca B. Mihalas, Sonali Arora, Samantha A. O’Connor, Heather M. Feldman, Christine E. Cucinotta, Kelly Mitchell, John Bassett, Dayoung Kim, Kang Jin, Pia Hoellerbauer, Jennifer Delegard, Melissa Ling, Wesley Jenkins, Megan Kufeld, Philip Corrin, Lucas Carter, Toshio Tsukiyama, Bruce Aronow, Christopher L. Plaisier, Anoop P. Patel, Patrick J. Paddison

## Abstract

In solid tumors, G0-like states are likely critical for maintaining developmental hierarchies and cellular heterogeneity and promoting tumor growth/recurrence, yet little is known about tumor G0 states or regulation of their ingress/egress. To discover G0-like states and their regulators for glioblastoma (GBM), we analyzed G0 populations in an orthotopic model of GBM using single cell RNA-seq and performed a genome-wide CRISPR-Cas9 screen in patient-derived GBM stem-like cells (GSCs) for genes that trap cells in G0 when inhibited. We identify the protein acetyltransferase KAT5 as a key regulator of transcriptional, epigenetic, and proliferative heterogeneity impacting transitions into G0-like states. KAT5 activity suppresses the emergence of non-dividing subpopulations with oligodendrocyte progenitor and radial glial cell characteristics both *in vitro* and in a human GSC brain tumor model. In primary gliomas, KAT5 activity is dynamic with KAT5^low^ tumor cells displaying quiescent properties, while KAT5 activity overall increases from low to high grade tumors and is associated with worse patient outcomes.

## Introduction

It is now well established from single cell RNA-seq studies that glioblastoma (GBM) tumors are complex, yet maligned, neuro-developmental ecosystems, harboring diverse tumor cell types, including cells resembling astrocytes, neural progenitors, oligodendrocyte progenitor cells, mesenchymal cells and radial glial cells, all of which presumably contribute to tumor growth and homeostasis in specific ways (e.g., ^1, 2^). Rather than deterministic hierarchies, this cellular and developmental heterogeneity may be better explained by phenotypic plasticity^3^, where cells can transition from one state to another through aberrant development paths not observed in healthy tissues.

However, single cell data sets have failed to produce general models for transitions in and out of specific developmental and proliferative states in tumors. One reason is that, for GBM tumor cells, there are no pre-existing *universal* markers that neatly resolve subpopulations into quiescent, “primed”, G1, or differentiated cellular states^4, 5^ as is the case, for example, for adult mammalian neurogenesis^6, 7^.

From histopathology studies, it is estimated that ~30% of glioblastoma tumor cells at a given moment are actively dividing (e.g., stain positive for the proliferation marker Ki67)^8^. The phenotypic behavior of the remaining ~70% of cells is largely unknown; although it is presumed that some portion are in a state of transient or long-term quiescence^4, 5^. The interplay between pathways and signals promoting cell cycle ingress and those governing its egress likely play key roles in generating cellular heterogeneity observed in GBM as well as its resistance to chemoradiation. Here, we sought to identify cellular states associated with G0-like states in human GBM stem-like cells (GSCs) as well as key regulators of these states.

## Results

### Single cell gene expression analysis reveals candidate G0-like populations

To functionally define GBM tumor cells in short- and long-term G0-like states, an orthotopic brain tumor derived from a human patient-derived GBM stem-like cells (GSCs)(GSC-0827) was used in conjunction with scRNA-seq and a reporter for G0 (p27-mVenus)(**Fig. 1a**; **Extended Fig. 1**). Because p27^hi^ cells have little or no G1 Cyclin/CDK activity, which is required for cell cycle reentry, the p27^hi^ population will be composed of cells entering G0-states (e.g., just after mitosis) and those already in short- or long-term quiescence. Thus, comparing scRNA-seq data of p27^hi^ to total tumor cells should reveal which cells in the tumor populations are more likely to be in G0-like states.

**Figure 1:**
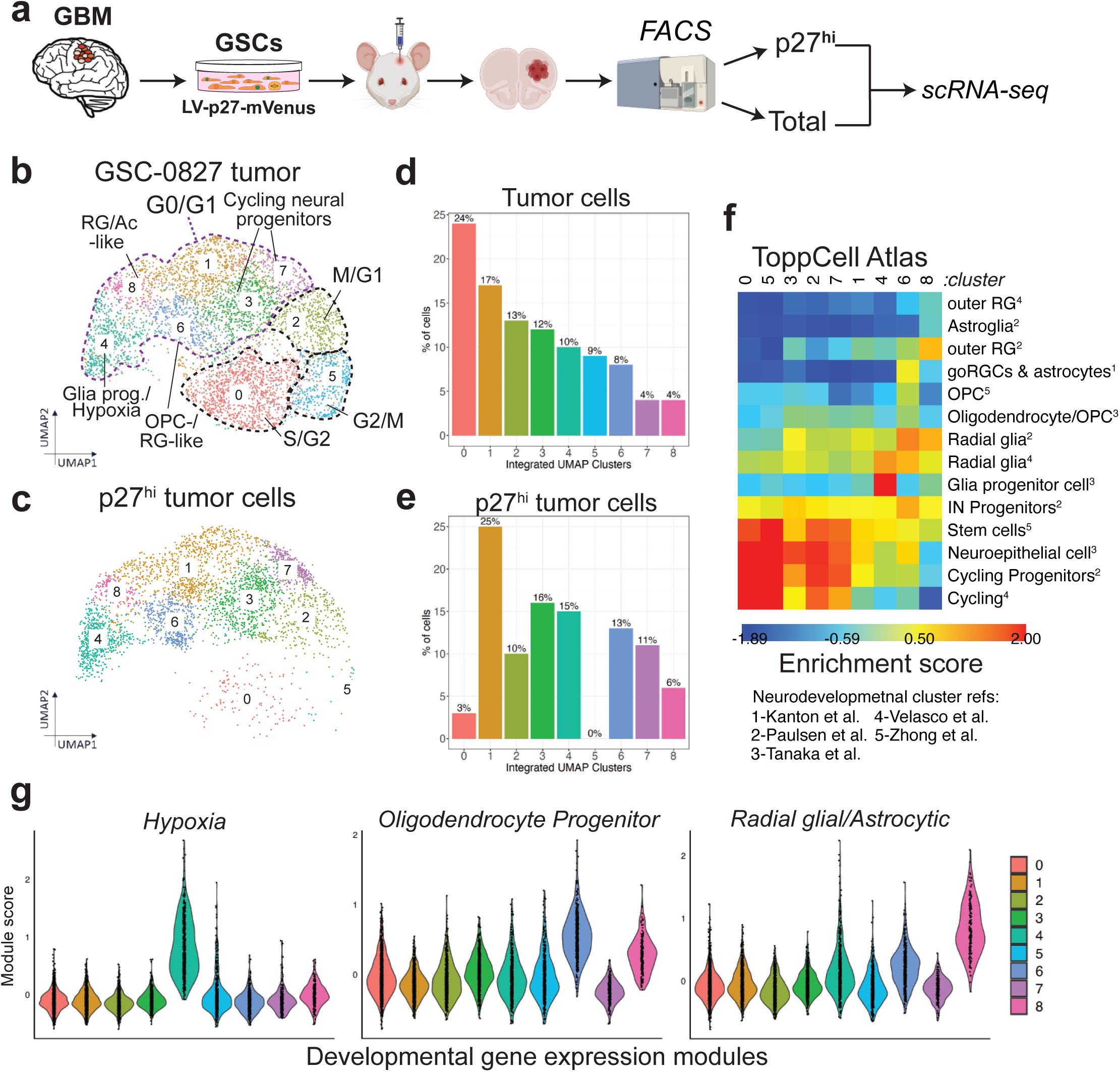
Single cell gene expression analysis of GSC-0827 tumors. **a**, Experimental overview for collection of samples for scRNA-seq. **b**, Projections of scRNA-seq data for GSC-0827 tumor reference using 3 tumor samples, along with inferred cluster cell cycle phase and neurodevelopmental associated cell type. GSC-0827 cells are an IDH1^wt^ proneural isolate^75, 84^. 5144 total tumor cells passed quality control metrics (Figure S2) and were used for downstream normalization and dimensional reduction using principal component analysis (PCA) and subsequent shared nearest neighbors clustering^85^. Data was visualized using uniform manifold approximation and projection (UMAP) for dimensional reduction of data and generation of *de novo* cell-based clusters^9^. Overview of experiment, filter cutoffs, and QC analysis are available in **Supplementary Fig. 1**. Supporting data includes: top enriched genes for each cluster (**Supplementary Table 1; Extended Data Fig. 2**); gene set enrichment (**Supplementary Tables 2-3**); gene expression modules and transcription factor networks (**Extended Data Fig. 3; Supplementary Tables 4-5**); and individual and gene set expression profiles (**Extended Data Fig. 4**). **c**, The tumor reference from **a** was used for mapping tumor cells from scRNA-seq analysis performed on GSC-0827 p27-mVenus^hi^ sorted tumor cells. 4780 p27^hi^ tumor cells passed quality control metrics. **d-e**, Proportion of cells in each cluster from **b-c**. **f**, Heatmap showing enrichment scores for scRNA-seq clusters in GSC-0827 tumor reference (A) for select neurodevelopmental cell types found in the ToppCell Atlas. A complete version of this figure is available in **Supplementary Fig. 2**. **g**, Select gene expression module scores for clusters in in GSC-0827 tumor reference. A list of genes in each module can be found in **Supplementary Table 4**. Each data point = single cell. Other gene modules and examples of expression of genes found in these modules are available in **Extended Data Fig. 3**.

Using uniform manifold approximation and projections (UMAP) for dimensional reduction of data and generation of *de novo* cell-based clusters^9^, we were able to identify 9 cell clusters in the GSC-0827 tumor reference (**Fig. 1b**) and their corresponding cell clusters in p27^hi^ cells (**Fig. 1c**), along with the relative frequencies of cells within each cluster (**Fig. 1d-e**).

To define cycling cell clusters, we employed differential gene expression (**Supplementary Table 1; Extended Data Fig. 2**), gene set enrichment analysis for each cluster (**Supplementary Tables 2-3**), and a cell cycle classifier (**Extended Data Fig. 3**). We identify three prominent clusters associated with cell cycle (0, 5, and 2). Cluster 0 is enriched in S-phase genes and cluster 5, G2/M genes (**Fig. 1b**). Both are conspicuously absent in p27^hi^ cells (**Fig. 1c-e**). Cluster 2 represents early G1/M transitional state where cells are in the process of exiting mitosis and still have residual mRNAs that peak in mitosis (**Extended Data Fig. 2**), which is also observed in hNSCs^5^. All other clusters showed significant reduction in cell cycle gene transcription (e.g., G1/S transition, S-phase, and mitotic genes), and metabolism gene sets (e.g., glycolysis, NAD/NADH metabolism, and oxidative phosphorylation) (**Supplementary Table 3**). We collectively designate these as G0/G1 (**Fig. 1b**), which are consistent expression patterns of individual cell cycle genes (**Extended Data Fig. 4**). From this analysis, GSC-0827 tumors are highly proliferative with ~33% of cells in S/G2/M, in line with measurements of Ki67 expression in primary GBM tumors (median = 27.5%)^8^.

To assess G0/G1 clusters, we used gene module comparisons for gene sets enriched in human neurodevelopment cell subtypes ^10^ (**Fig. 1f**) and previously proposed GBM cellular subtypes (e.g., ^2^) (**Fig. 1g**). This analysis revealed G0/G1 clusters prominently associated with oligodendrocyte progenitor cells (cluster 6) (OPC), radial glial cells/astrocytes (RG) (cluster 8), cycling neural progenitors (clusters 3/7), and hypoxia (cluster 4).

To reveal cellular transitions from mitosis to G0-like states, we performed mRNA velocity analysis, which models the direction and speed of cell dynamics in scRNA-seq data based on kinetics of transcription and splicing^11^ (**Fig. 2a-b**). For cycling cells, this analysis revealed that tumor cells reenter the cell cycle from clusters 2 (M/G1), 3 (cycling progenitors), or 6 (OPC). However, cluster 1 cells (associated with cycling neural progenitors (**Fig. 1f**)) also show expression of the G1/S cyclin, CCNE2, DNA replication genes expression (G1/S) (**Extended Data Fig. 4**), and E2F1 expression (**Fig. 2c-e**). The latter is part of a sequenced expression of cell cycle transcription factors, which includes MYBL2, FOXM1, and MYC, each of which promotes phase-specific cell cycle gene expression, starting with E2F1^12^ (**Fig. 2c-e**).

**Figure 2:**
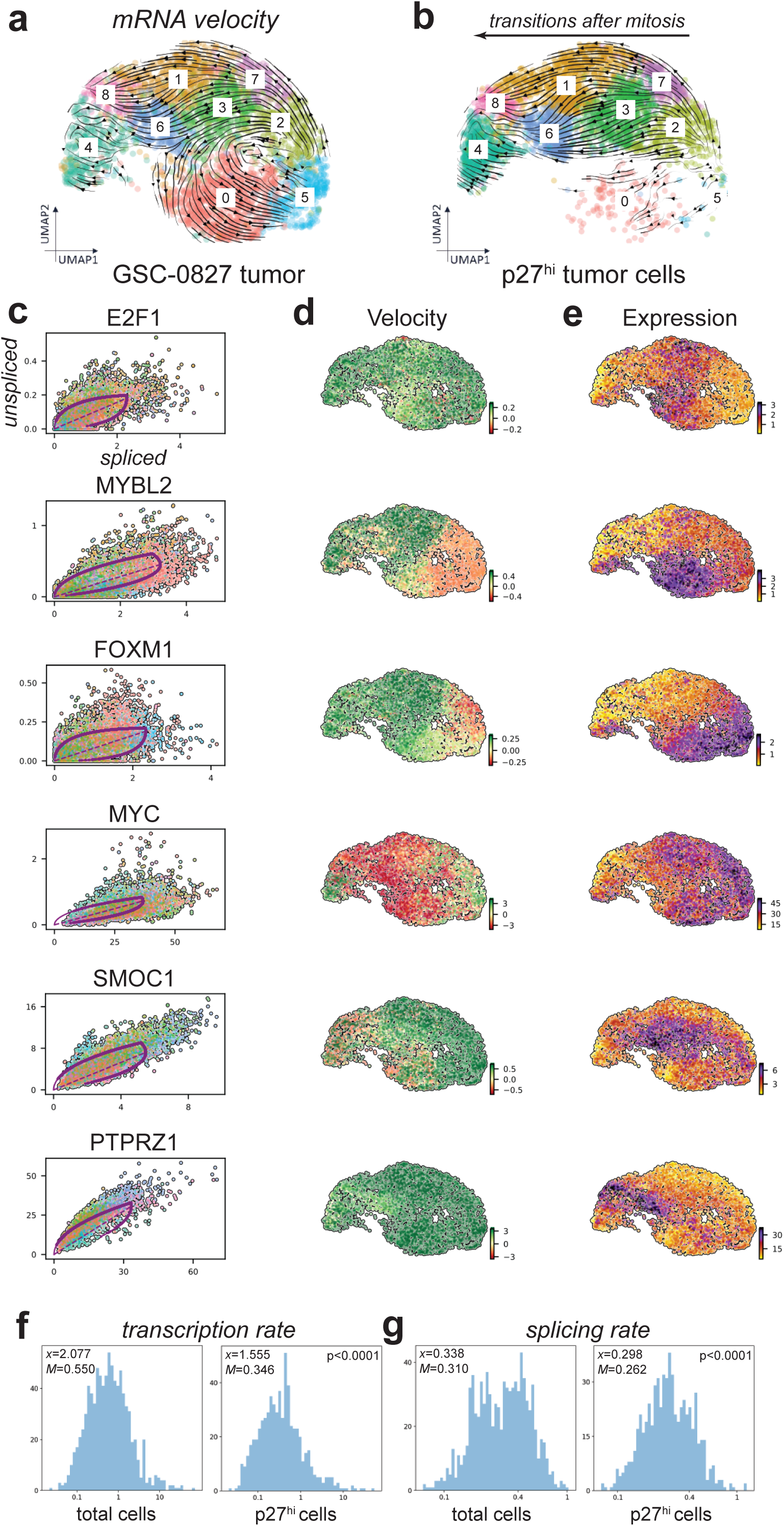
mRNA velocity analysis of total and p27hi cell populations in GSC-0827 tumors. **a-b**, RNA velocity analysis of scRNA-seq data for Figure 1a-b. **c-e**, Key cell cycle transcription factors with significant mRNA velocity differences in GSC-0827 scRNA data. **c**, This column of graphs shows unspliced versus spliced ratio of candidate genes with significant changes in velocity between clusters, where each data point is a cell colored by cluster from **a**. Contoured lines are derived from dynamical modeling of mRNA velocity from scVelo (https://scvelo.readthedocs.io/) (Bergen et al., 2020). Each of these genes are “driver genes” for velocity changes and lines for one or more clusters (Bergen et al., 2020). A complete list is available in **Supplementary Table 6**. **d,** Velocity values for each gene mapped onto UMAP projection of GSC-0827 tumor reference. High velocity values for a particular gene (i.e., higher predicted unspliced/spliced mRNA ratios) tend to occur before expression peaks, followed by lower velocity and transcriptional down regulation. **e**, Gene expression values for each gene mapped onto UMAP projection of GSC-0827 tumor reference. **f-g**, Histograms of estimated transcription rates and splicing rates for genes for which scVelo dynamic models of transcription and splicing are available (n=1032 genes for total tumor cells; n=749 for p27^hi^ cells)(*x*=mean; *M*= median). A two sample Kolmogorov–Smirnov test was used to test significance.

RNA velocity analysis revealed that no p27^hi^ cells reenter the cell cycle (**Fig. 2b**). Instead, cells transition from M/G1 along two general tracks, either through cluster 3 to 6 (OPC) and then 8 (RG/Ac) or through clusters 3 or 7 to 1 and then 8. Of note, cells along the 7-1-8 cluster track express CD44 (a marker of mesenchymal GBM cells) and appear largely mutually exclusive from cells that express OLIG1 (an OPC marker) (**Extended Data Fig. 4**). However, both trajectories terminate in RG-like cluster 8. Because we know where cells exit the cell cycle, the data support a model whereby tumor cells with longer resident times in G0-like states ultimately adopt OPC-like or RG-like transcriptional features.

Cross comparisons of genes enriched in clusters 6 and 8 revealed overlap of key OPC and RG/Ac-associated genes and transcription factors (**Extended Data Fig. 3**). These included, for example: <u>OPC</u>: ASCL1, BCAN*, OLIG1*, NFIA, PLP1, PTPRZ1*, SOX6; <u>RG/Ac</u>: CLU*, ID1/3/4, F3, GFAP, LZTS1, PTN*, SPARC*, SLC1A3, TTYH1. [*Genes among top five differentially expressed for these clusters]. Many of these genes participate in neurogenesis or glioma biology by promoting stemness, proliferation, invasive behavior, key signaling pathways (e.g., Notch), or survival in GSCs and/or NSCs (referenced in ^5^). p27^hi^ populations also in general displayed lower rates of transcription and splicing (**Fig. 2f-g**). Lower transcription rates and RNA content are hallmarks of quiescent cells^13, 14^.

Of note, cluster 4 (glia progenitors/hypoxia) presents as separate G0-like state. This cluster shows prominent and almost exclusive expression of multiple hypoxia-associated genes including NRDG1 and VEGFA^15, 16^ (**Extended Data Fig. 2 and 4**). However, because there are few velocity lines leading there, it is difficult to assess whether cells transition from normoxia to hypoxia after M/G1 or from other states or both.

### A functional genomic screen to identify genes that regulate G0-like states in human GSCs

Taken together, the above results reveal tumor cell states associated with transient and, likely, longer term quiescent populations in GSC-0827 tumors. We next wished to identify key regulators of G0 ingress/egress in these same tumors. To this end, we used GSC-0827 p27 reporter cells in a genome-wide knockout screen to identify genes which when inhibited trap cells in G0-like cells (**Fig. 3a**). Comparing sgRNAs in unsorted versus sorted double positive populations yielded 75 genes enriched and 37 depleted in p27^hi^CDT1+ cells (FDR<.01) (**Fig. 3b**). The most prominent enriched gene sets included those involved in ribosome assembly and ribosome protein coding genes (e.g., RPL5, RPS16,) and the Tip60/NuA4 lysine acetyltransferase complex (e.g., ACTL6A, EP400, KAT5, TRAPP), with the KAT5 catalytic subunit scoring among top ten screen hits (**Fig. 3b; Extended Fig. 5**; **Supplementary Table 7**).

**Figure 3:**
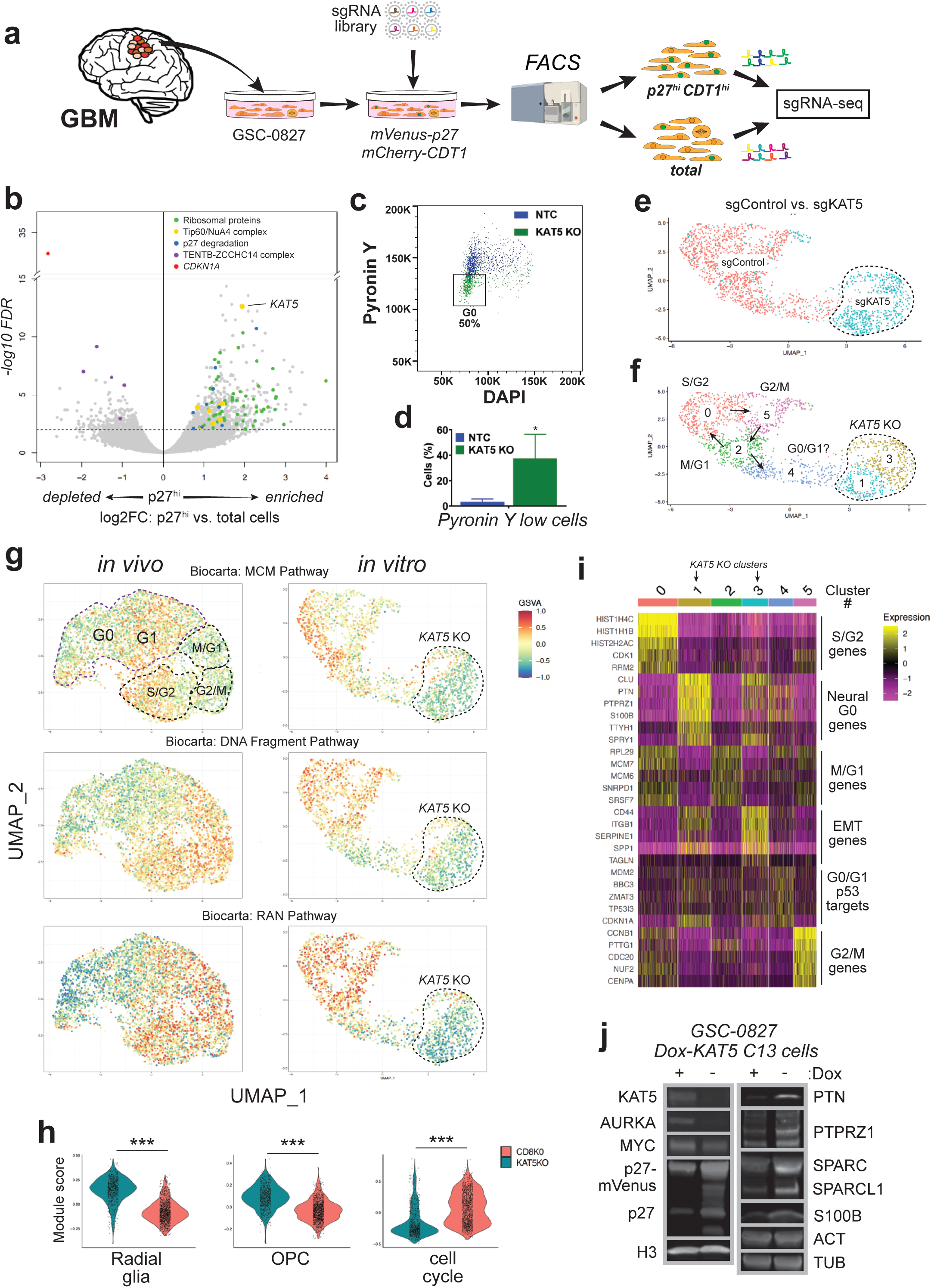
Identification of KAT5 as a G0-trap in GSC-0827 cells. **a**, Schematic of the G0-trap screen. For the screen, GSC-0827 cells containing mCherry-CDT1 (G0/G1) and p27-mVenus (G0) were transduced with a genome-wide CRISPR-Cas9 library, allowed to expand for 10 days, and sorted for double positive cells using the top 20% of p27-mVenus+ cells as a cut off. **b**, Results G0-trap screen from sgRNA-seq of p27^hi^ vs. total cell population (n=3; edgeR was used to assess p values and logFC cutoffs) (**Supplementary Table 7**). Supporting QC data and gene set enrichment can be found in **Extended Data Fig. 5**. **c**, FACS-based assessment of total RNA and DNA content, using pyronin Y and DAPI, respectively, in GSC-0827 cells via nucleofection of sgRNA:Cas9 RNPs. Additional retest assays are available in Extended Data Fig. 6 and 7. **d**, Quantification of **c** (n=3; student’s t-test, p<.01). **e**, UMAP projection of scRNA-seq data for sgCD8 (control) and sgKAT5 cells in GSC-0827 cells 5 days post nucleofection. Filtering scheme and data quality assessment for this data is available in **Supplementary Figure S3**. **f**, The UMAP projection from **e** showing *de novo* clusters generated. Cell cycle state predictions using the ccSeurat and ccAF classifiers and RNA velocity analysis for this data from are available in **Extended Data Fig. 8**. Associated data files include: cluster-based gene expression analysis (**Supplementary Table 8**) and gene set enrichment for top 200 expressed and top 200 depleted genes for each cluster (**Supplementary Tables 9 and 10**, respectively). For control cells, there is one G0-like cluster (cluster 4) which showed some weak expression for p53-associated target genes, suggestive of a DNA damage-induced G0-like state, a common feature of cultured cells ^17, 86^ also to some degree Neural G0 genes (see text). This state likely represents the p27^hi^ Edu-population observed *in vitro* in **Extended Data Fig. 1b**, which are capable of re-entering the cell cycle. **g**, Gene Set Variation Analysis (GSVA) associated with clusters and cell cycle phases for GSC-0827 tumor reference (Fig. 1b) and *in vitro* KO of KAT5 from **f**. **h**, Violin plots of gene expression module scores for each cell from scRNA-seq data of sgCD8A and sgKAT5KO GSC-0827 cells. oRG: outer radial glia. OPC: oligodendrocyte precursor cells. MES: mesenchymal. Each data point is a single cell. Wilcoxon signed ranked test and t-tests were used to assess significance (p<.001 for both). Genes contained in each module are available in **Supplementary Table 4**. **i**, Heatmap of representative genes upregulated in scRNA-seq clusters from **f**. **j**, Western blot validation studies of gene expression changes associated with loss of KAT5 activity. Protein extracts from GSC-0827 C13 cells were used from Dox+ or Dox- (7 days) conditions.

Among sgRNA targets significantly depleted in p27^hi^CDT1+ cells, we find CDKN1A/p21 as a top scoring hit. CDKN1A/p21 is a cyclin/CDK2 inhibitor that has been shown to promote entry into G0-like states in cultured cells^17^, consistent with the screen results. In addition, we find genes associated with the TENTB-ZCCHC14 complex including PAPD5 and ZCCHC14, which help destabilize rRNAs and other non-protein coding RNAs, including miRNAs and TERC^18–20^.

Retests of several enriched and depleted G0-trap screen genes largely recapitulated the screen data (**Extended Fig. 6a-b**). KO of RPS16, RPL5, and KAT5 complex members, resulted in significant increases in p27-mVenus^hi^ subpopulations and loss of proliferation in GSC-0827 cells, while KO of CDKN1A and PAPD5 decreased steady-state p27 levels and increased proliferation (**Extended Fig. 6b-c**). However, RPS16 and RPL5 KO resulted in significant cell toxicity, while KAT5 KO cell numbers appeared to be stable even for extended periods (e.g., >2 weeks) (see below).

That genes involved in ribosome function would score as G0 trap mutants is supported by the notion that down regulation of protein synthesis and ribosome assembly are hallmarks of quiescent cells^21, 22^; and, conversely, that their activity increases as, for example, neural stem cells transition out of quiescence into an activated state that precedes cell cycle entry^23^.

The KAT5/NuA4 lysine acetyltransferase complex targets both histones (H2A variants, H3, and H4) and non-histone proteins for acetylation and functions as a transcriptional co-activator, whose activities are coordinated with multiple transcription factors ^24–26^. In addition, the NuA4 complex also participates in DNA double-strand break repair by facilitating chromatin opening^27, 28^. However, little is known about the role of KAT5/NuA4 complex activity in GBM biology. Therefore, we chose to further pursue the question of whether KAT5 activity affects transitions in and out of G0-like states in GBM cells.

### KAT5 knockout triggers a G0-like state in *in vitro* cultured human GSCs

We examined multiple phenotypes associated with G0-like states using the nucleofection KO assay. First, we determined whether KAT5 KO could induce a population of cells with G1 DNA content and low total RNA content, a classic indicator of quiescent cells. Five days after introduction of sgKAT5:Cas9 complexes, a significant G1/G0 RNA^low^ population emerges in GSC-0827 cells (**Fig. 3c-d**). Analysis of cell cycle proportions via FUCCI factors (**Extended Fig. 6e-g**) and DNA synthesis rate via EdU incorporation (**Extended Fig. 7a-b**) were also consistent with this result. In each case, KAT5 KO significantly altered these measures in GSC-0827 cells, reducing the frequency of Geminin^hi^ and EdU^hi^ cells, respectively. Similar results were also observed in five additional human GSC isolates (**Extended Fig. 7c**) and also showed upregulation of p27 (**Extended Fig. 7d**).

Another key hallmark of G0-like states is reversibility, which distinguishes quiescence from differentiation and senescence. Adding back KAT5, 7 days after KAT5 KO caused the same rapid accumulation of cells as parental controls and the return of parental ratios of cell cycle phases (**Extended Fig. 7e-g**). Taken together, above results are consistent with KAT5 activity controlling reversible ingress/egress into G0-like states in GSCs.

### Single cell gene expression analysis of KAT5 inhibition in GSC-0827 cells *in vitro*

We next investigated cellular states induced by KAT5 inhibition in cultured GSCs *in vitro* by performing scRNA-seq analysis in GSC-0827 cells (**Fig. 3e-f**; **Extended Fig. 8a-c**). By this analysis, control GSC-0827 cultures have an embryonic stem cell-like cell cycle, with ~80% of cells in three cycling cell clusters: M/G1 (cluster 2), S/G2 (cluster 0), and G2/M (cluster 5). This is in contrast to *in vivo* results whereby ~45% of tumor cells are in these phases (side by side UMAP projections are shown in **Fig. 3g**).

However, KAT5 KO causes the cell cycle clusters and cell cycle gene expression to collapse and two new clusters to emerge (clusters 1 and 3). These new states have significant increases in OPC and radial glial gene expression and loss of cell cycle and nuclear pore gene transcription, more similar to G0 states found *in vivo* (**Fig. 3h-i**). These include “Neural G0” genes^5^, which include many genes prominently up regulated in tumor clusters 6 (OPC) and 8 (RG/Ac) and are known to participate in neurogenesis or glioma biology. Some of these include: CLU, a secreted antiapoptotic factor^29^; F3, a marker of quiescent GBM cells^30^; PTN and its target PTPRZ1, which promote stemness, signaling, and proliferation of neural progenitors and glioma tumor cells^31–34^; S100B, a chemoattractant for tumor-associated macrophages in glioma^35, 36^; and SPARC and its ortholog SPARCL1, which promote brain tumor invasion and survival^37–39^; and TTYH1, required for NSC stemness and Notch activation^40, 41^. Western blot analysis confirmed up regulation of many of these genes after KAT5 inhibition in GSC-0827 cells, along with down regulation of cell cycle-associated proteins AURKA and MYC and stabilization of endogenous p27 (**Fig. 3j**). F3 upregulation was confirmed by FACS analysis (**Extended Fig. 8d**).

In addition, Cluster 3 genes up regulated show enrichment for interactions with the extracellular matrix, integrin signaling, and cell migration (**Supplementary Table 9**), which includes genes that participate in the proneural-to-mesenchymal transition in GBM tumors (e.g., CD44, CDH2/N-Cadherin, SERPINE1, SPP1/Osteopontin)^42^ (However, this was not observed *in vivo*, below). Taken together, these data are consistent with KAT5 KO triggering a G0-like state not otherwise apparent *in vitro* in GSC-0827.

### KAT5 loss in GSC-0827 cells *in vivo* alters sites of mixed epigenetic valency to promote emergence of G0-like populations in GSC-0827 tumors

To determine *in vivo* responses to KAT5 loss, we used GSC-0827 cells that were engineered to have a doxycycline (Dox) controllable KAT5 expression (C13 cells) (**Fig. 4a**). For C13 cells, Dox withdrawal (KAT5^off^) caused loss of “pan” H4-acetylation (ac) levels and also monoacetylation at K12 (a key target of KAT5^43^), while total post-translational marking of histone H3 or total histone levels remained unaffected (**Fig. 4b**). Over the course of 7 days, KAT5^off^ triggered a progressive induction of p27 in cells and concomitant lowering of H4-Ac in the same cells (**Fig. 4c**).

**Figure 4:**
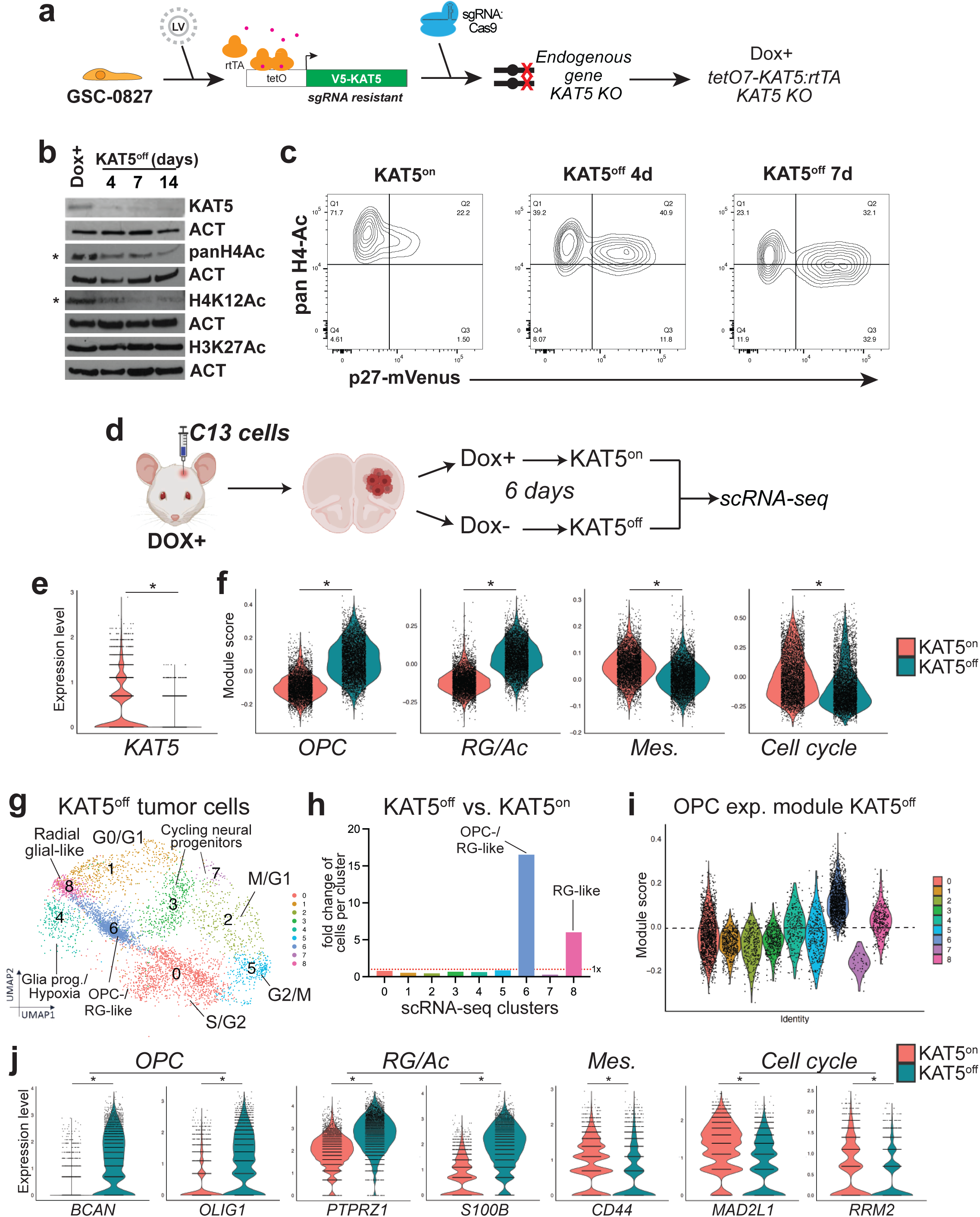
Single cell gene expression analysis of GSC-0827 tumors after KAT5 inhibition *in vivo*. **a**, GSC-0827 cells were engineered to have a doxycycline controllable KAT5 open reading frame and knockout insertion-deletion mutations in the endogenous KAT5 gene. We found one such clone, #13 (C13), to have Dox-dependent KAT5 expression and requirement of Dox+ for continued growth. **b**, Western blot analysis of KAT5 and histones H3 and H4 acetylation and methylation status after Dox withdrawal of 0, 4, 7, and 14 days in C13 cells. *indicates KAT5 targets. **c**, FACS analysis of histone H4 acetylation and p27-mVenus levels after 0, 7, and 14 days of Dox withdrawal in C13 cells. **d**, Overview of *in vivo* tumor experiments with C13 cells. After MRI confirmation of tumor formation, Dox was either maintained or was removed from drinking water for 6 additional days. **e**, Gene expression changes for KAT5 in KAT5^on^ vs. KAT5^off^ C13 tumors. **f**, Expression model scores for KAT5^on^ vs. KAT5^off^ C13 tumors. **g**, UMAP projection of scRNA-seq data for C13 KAT5^off^ cells using 3 tumor samples, along with inferred cluster cell cycle phase and neurodevelopmental associated cell type. **h**, Fold change of cells in each scRNA-seq cluster for KAT5^off^ tumors. **i**, OPC expression module scores for clusters in tumor clusters for KAT5^off^ tumors. **j**, Violin plots of gene expression module scores for each cell from scRNA-seq data of Dox+ vs. Dox-GSC-0827 C13 tumors. Each data point = single cell. Wilcoxon signed ranked test and t-tests were used to assess significance (p<.001 for both). Genes associated with each model are available in **Supplementary Table 4**.

We then used C13 cells to generate GBM tumors in immunocompromised mice. In this scenario, mice are administered Dox in their drinking water prior to injection to support engraftment of C13 cells. After confirmation of tumor formation by MRI (~3 weeks), Dox was either maintained (KAT5^on^) or removed for 6 days (KAT5^off^), after which scRNA-seq was performed (**Fig. 4d**).

Consistent with manipulating KAT5 gene expression, the KAT5^off^ state results in significant reduction in the KAT5 transgene (**Fig. 4e**). KAT5^off^ shows significant increases in OPC and RG-cluster associated gene expression and concomitant with reductions in mesenchymal and cell cycle gene transcription compared to KAT5^on^ (**Fig. 4f**). Consistent with this result, we observed increases in the relative KAT5^off^ cell numbers in G0-like OPC and RG-like clusters (**Fig. 4g-i**), resulting in a ~16-fold increase in cells in OPC-like cluster, representing 33% of the overall tumor cells and a 6-fold increase in the RG-like cells, while other G0/G1 populations decreased (**Fig. 4h**). Examples of KAT5^on^ vs. KAT5^off^ changes in expression of genes from OPC, RG/Ac, mesenchymal, and cell cycle gene modules are shown in **Fig. 4j**.

Next, to understand chromatin state changes occurring in KAT5^on^ vs. KAT5^off^ tumor cells, we performed CUT&TAG analysis^44^ for activating H3K4me2 and H3K27ac marks, found at both promoters and enhancers^45^, and repressive H3K27me3 marks, required for long-term epigenetic stability of cell states during normal differentiation^46, 47^.

Chromatin state discovery analysis (i.e., ChromHMM)^48^, which integrates the chromatin marks and generate layered “emission states” across each region of the genome, identified 8 emission states in GSC-0827 KAT5^on^ tumors (**Fig. 5a**). These included two “mixed valency” states, which contain chromatin regions with both activating and repressive marks: E4, which contains all three marks, and E6, which only contains H3K4me2 and H3K27me3. These regions likely reflect tumor clusters with different transcriptional states (i.e., active in one subpopulation and repressed in another).

**Figure 5:**
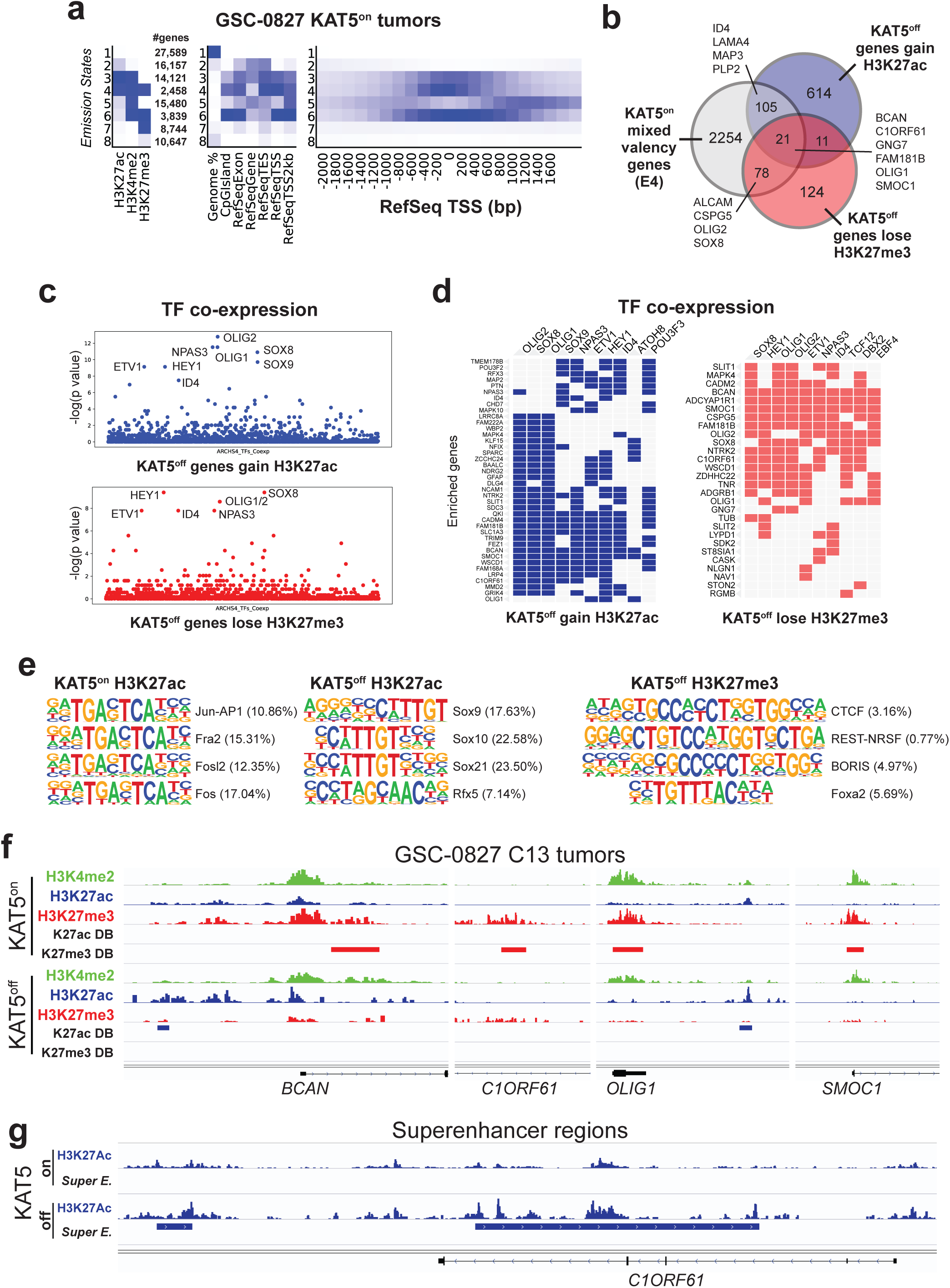
Analysis of epigenetic patterning associated with KAT5^on^ and KAT5^off^ states in GSC-0827 tumors. **a**, ChromHMM analysis of genomic regions in KAT5^on^ C13 tumor cells showing 8 possible chromatin states (i.e., emission states) for H3K4me2, H3K27ac, and H3K27me3, the associated number of genes, and emission state region genomic annotations. Genome%= intergenic space; TES = transcription end sites; TSS = transcription start sites. The darker blue color corresponds to a greater probability of observing the mark in the state. The full data set is available in **Supplementary Table 11**. **b**, Overlap of genes associated with emission state E4 from **a**, which display both activating and repressive chromatin marks, and those with significant changes H3K27ac and H3K27me3 after loss of KAT5 activity in GSC-0827 tumors. DiffBind and DESeq2 were used to score significant changes in chromatin marks (**Supplementary Tables 12 and 13**) from KAT5^on^ and KAT5^off^ C13 tumor samples shown in Figure 4d-e (n= 2). Note: DiffBind could not be performed on H3K4me3 marks for C13 tumor samples because one replicate failed to produce sufficient quality data. **c**, Enrichment for genes associated with expression of human transcription factors (TFs). Top panel: genes enriched for H3K27ac+ regions in after loss of KAT5 activity. Bottom panel: genes enriched for H3K27ac+ regions in after loss of KAT5 activity. The analysis was performed using the Enrichr pipeline (Kelshov et al., 2016) using the hypergeometric test by comparing the top 300 genes associated with specific human transcription factors in the ARCHS4 database (Lachmann, et al., 2018). **d**, Gene-based output from **c** using colored squares indicate genes enriched in specific TF associated gene sets from ARCHS4 database. **e**, Enrichment for known transcription factor binding sites associated with H3K27ac and H3K27me3 sites are enriched in KAT5^on^ and KAT5^off^ C13 tumors. Dox+ condition for H3K27me3 did not yield significant enrichments. The complete datasets can be found in **Supplementary Table 14**. **f**, Examples of multivalent genes in KAT5^on^ C13 tumors that upon Dox withdrawal gain H3K27ac and lose H3K27me3 marks (i.e., overlapping genes from **b**). Additional examples including GNG7, SLIT1, and SOX8 are shown in **Extended Data Fig. 9b-c**. g, An example of the superenhancer site for C1ORF61 showing H3K27Ac tracks for KAT5^on^ and KAT5^off^ tumors along with called S.E. site. Additional superenhancer callouts KAT5^on^ and KAT5^off^ can be found in **Extended Data Fig. 9d-e**. The complete data sets can be found in **Supplementary Table 15**.

Both E4 and E6 had the fewest number of associated genes (2458 and 3839, respectively) and shared overlap of 1002 region-associated genes. Their regions have a higher probability of occurring in CpG islands and transcriptional start site of genes, for example, in contrast to regions with only H3K27me3 sites (**Fig. 5a**).

E4 associated genes had substantial overlap with H3K27ac and H3K27me3 genic regions significantly altered in KAT5^off^ tumors (**Fig. 5b**). These included many of the key genes associated with OPC and RG/Ac clusters in GSC-0827 tumors, including, for example, OLIG1 and OLIG2 (**Fig. 5b**), which lose H3K27me3 repressive marks and/or gain H3K27ac marks in KAT5^off^ tumor samples (**Supplementary Tables 12-13**). Most (~47%) of the H3K27me3 marks lost after Dox withdrawal occur within 1kb of promoter region of these genes (compared to only ~10% for total peaks) (**Extended Fig. 9a**).

There were multiple transcription factors associated with “activated” genes in KAT5^off^ tumors (i.e., gain of H3K27ac and/or loss of H3K27me3) (**Fig. 5c-d**). These included key neurodevelopmental transcription factors expressed in NSCs (ID4, SOX9), astrocytes (HEY1, ID4, and SOX9) or oligodendrocyte lineages (OLIG1, OLIG2, and SOX8) in normal brains and also glioma^49–51^, and also NPAS3 which has been associated with glioma indolence^52^.

Further, >20% of these regions with H3K27ac peaks that increase in KAT5^off^ tumors show enrichment for the degenerate Sox transcription motif (i.e., CWTTGT) (**Fig. 5e**; **Supplementary Table 14**), which is required for Sox transcription factor binding, transcriptional activity, and neurodevelopmental cell patterning/function^53^. By contrast, H3K27ac peaks enriched in KAT5^on^ tumors contained sites bound by AP-1 transcription complex members, which play key roles in promoting mesenchymal gene expression in multiple cancers, including GBM^54, 55^ (**Fig. 5e**).

Callouts for multiple gene loci affected by KAT5^off^ are shown in **Fig. 5f** and **Extended Fig. 9b-c**. These include genes that show mixed valency in KAT5^on^ tumors, including key regulators of OPC lineage identity, including OLIG1 and SOX8, as well as genes associated with OPC and RG/Ac GSC-0827 tumor clusters, such as BCAN, C1ORF61, GNG7, SLIT1, and SMOC1. Many of these genes show specific expression in normal human OPCs (www.proteinatlas.org).

In addition, we investigated KAT5 driven changes in superenhancers, i.e., multi-enhancer regions (<12.5kb) defined by H3K27ac marks^56^, in tumor samples (**Fig. 5g; Extended Fig. 9d-e**; **Supplementary Table 15**). For KAT5^on^ tumors, we identified and ranked 521 superenhancer regions associated with genes implicated in GBM mesenchymal cell states (e.g., BCL3, CD44, RGS16), GBM survival (SEC61G, VGF), neurogenesis/stemness (PBX1), and cell cycle (CDK6) (**Extended Fig. 9d**). For KAT5^off^ tumors, 218 superenchancer regions were associated with many of the OPC and RG/Ac genes highlighted above, including BCAN, C1ORF61, GNG7, SMOC1, and SOX8, as well others including angiotensinogen (AGT), a biomarker for bevacizumab response in GBM, CSPG5, a marker of OPC cells, DBI, associated with quiescent cancer cells, and GFAP, a key astrocytic marker found in the RG/Ac cluster (**Fig. 5g; Extended Fig. 9e**).

### Attenuating KAT5 activity in GSCs shifts cell cycle and growth dynamics towards a G0-like state *in vitro* and *in vivo*

The above results show that after complete loss of KAT5 GBM cells enter a G0-like state and detail many of the accompanying phenotypic, transcriptional, and epigenetic changes. We next wished to determine if partial attenuation of KAT5 activity would affect GSC growth *in vitro* and tumor growth *in vivo*.

To this end, we first performed Dox titration experiment with cultured C13 cells. Growing C13 cells in Dox concentrations between 0, 25, 50, 100, and 1000 ng/mL slowed growth rates progressively increased p27-mVenus reporter activity (**Fig. 6a-b**). At the same time, significant and progressive attenuation H4-Ac was also observed (**Fig. 6c**). Thus, incremental reduction in KAT5 expression in C13 cells leads to incremental loss of KAT5 activity, cell growth, and increases in G0 populations of cells.

**Figure 6:**
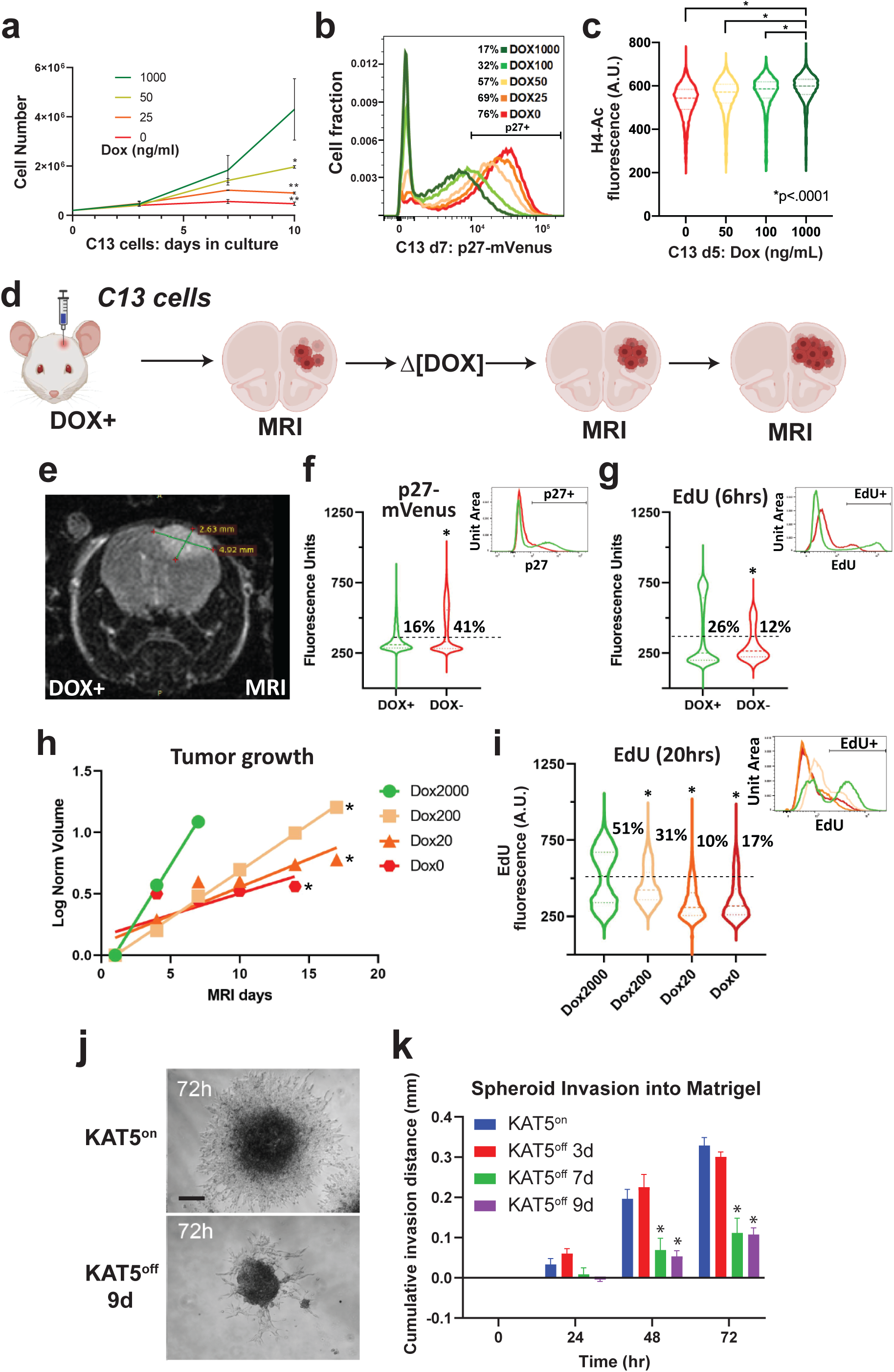
Modulation of KAT5 activity in GSC-derived tumors and during GSC invasion assays. **a**, A growth curve with C13 cells grown in various concentrations of Dox (n=3; student’s t-test, p<.05). **b**, FACS analysis of p27 induction in C13 cells grown in shown concentrations of Dox for 7 days. **c**, Violin plot of H4-Ac levels in C13 cells grown in shown concentrations of Dox for 5 days. KS test was used to test significance. *pval <.0001. **d**, Scheme for using C13 GSC-0827 cells for creating orthotopic xenograft tumors in NSG mice. **e**, Representative MRI image of mouse head with Dox-KAT5 tumor in cortex at 36 days post-injection. **f**, Analysis of p27 levels after Dox withdrawal in C13 tumor containing xenograft mouse brain. KS test was used to test significance (p<.0001). **g**, Analysis of EdU incorporation (6hrs) after 7 days of Dox withdrawal in C13 tumor containing mouse. KS test was used to test significance (p<.0001). **h**, Tumor growth as assessed by volume using MRI in C13-induced brain xenografts after switching drinking water to concentration of Dox indicated (µg/mL). Linear regression analysis was used to assess significance (p-val overall=0.001; p-val2000 vs. 200<0.0001; p-val2000 vs. 20=0.0049; p-val2000 vs. 0=0.0183). **i**, EdU incorporation for 20hrs at 14 days after Dox concentration from (E). KS test was used to test significance (p<.0001). **j-k**, KAT5^on^ vs. KAT5^off^ C13 invasion assays. C13 cells were grown as spheres for 3 days in either 1μg/ml Doxycycline for KAT5^on^ or no Doxycycline for KAT5^off^ conditions and transferred to Matrigel covered wells for the specified number of days (3d, 7d, or 9d). Phase-contrast images were captured every 24 hours for 72 hours. The area covered by invading cells was measured using FIJI. **j**, Representative phase contrast microscopy images of invasion assay. Spheroids were embedded into Matrigel. Scale bar is 200 μm. **k**, Quantification of (E). n_KAT5on_=3, n_KAT5off 3d_=3, n_KAT5off 7d_=4, n_KAT5off 9d_=5. *indicates p≤0.02, student’s t-test.

Next, we used C13 cells to generate GBM tumors (**Fig. 6d**). After confirmation of tumor formation by MRI (e.g., **Fig. 6e**), Dox was either removed or titrated. Dox withdrawal for 7 days resulted in significant induction in p27 and loss of Edu^+^ cells (**Fig. 6f-g**). Titrating Dox (0, 20, 200 and 2000 μg/ml) resulted in significant alteration of tumor growth (**Fig. 6h**). In full dose Dox, C13 tumors growth was similar to parental tumors (~43-48 day survival). The 200 μg/ml dose resulted in intermediate tumor growth and intermediate loss of EdU incorporation, while 20 and 0 μg/ml concentrations did not significantly differ from each other (Fig. 6h). The 200, 20, and 0 μg/ml Dox trials showed concomitant reductions in EdU incorporation (**Fig. 6i**). Taken tother, the results demonstrate that loss of KAT5 reduces tumor growth in pre-formed tumors.

However, a lingering question from these studies was whether KAT5^off^ cells would become more invasive as a result of entering G0-like states. We found the opposite was true: KAT5^off^ show significantly less invasiveness than KAT5^on^ cells (**Figs. 6j-k**).

### KAT5 activity in primary glioma cells

We next determined whether primary gliomas harbor populations of KAT5^low^ cells that have molecular features consistent with G0-like states. Previous work has established that low grade gliomas (LGGs) exhibit more indolent growth compared to high grade gliomas (HGGs)^57^ and almost universally harbor IDH1/2 mutations^58^. Based on gene expression analysis, LGGs have a higher proportion of G0-like cells compared to HGGs^5^. Although LGGs result in better responses to standard of care and extended survival times, they inevitably recur as HGG with IDH1/2 mut^59^. We performed cell-based analysis of freshly resected low- and high-grade primary gliomas. We took advantage of our FACS assay for KAT5 activity, which uses H4-Ac as a readout. We also chose to use protein synthesis rate as a secondary read out in these assays.

Because KAT5 loss leads to a significant reduction in total RNA levels (which is mainly comprised of rRNA), it is likely, we reasoned, that KAT5 controls rRNA transcription and, as a result, protein synthesis rate in GSCs. This was indeed the case. After KAT5 loss, there were significant reductions in rRNA levels and EU incorporation (which in mostly incorporated into rRNA) in KAT5^off^ C13 cells (**Extended Fig. 10a-b**). p27^hi^ cells in KAT5^on^ cells have both low EU incorporation rates and lower H4-Ac levels, while loss of KAT5 causes further reduction of both (**Fig. 4c**; **Extended Fig. 10b**).

Protein synthesis (assayed via L-azidohomoalaine (AHA) incorporation) was strongly suppressed after 5 days KAT5^off^ (**Fig. 7a**). However, GSC cells, none-the-less, retain the same cellular mass as control cells, even after 14 days of KAT5^off^ (**Fig. 7b**). With titration of Dox, there was progressive loss of protein synthesis in line with H4-Ac loss in these cells (**Fig. 7c**). Thus, KAT5 activity is linked to protein translation rates in these cells and attenuating KAT5 activity likely triggers slower G0 egress, in part, to conserve cellular mass.

**Figure 7:**
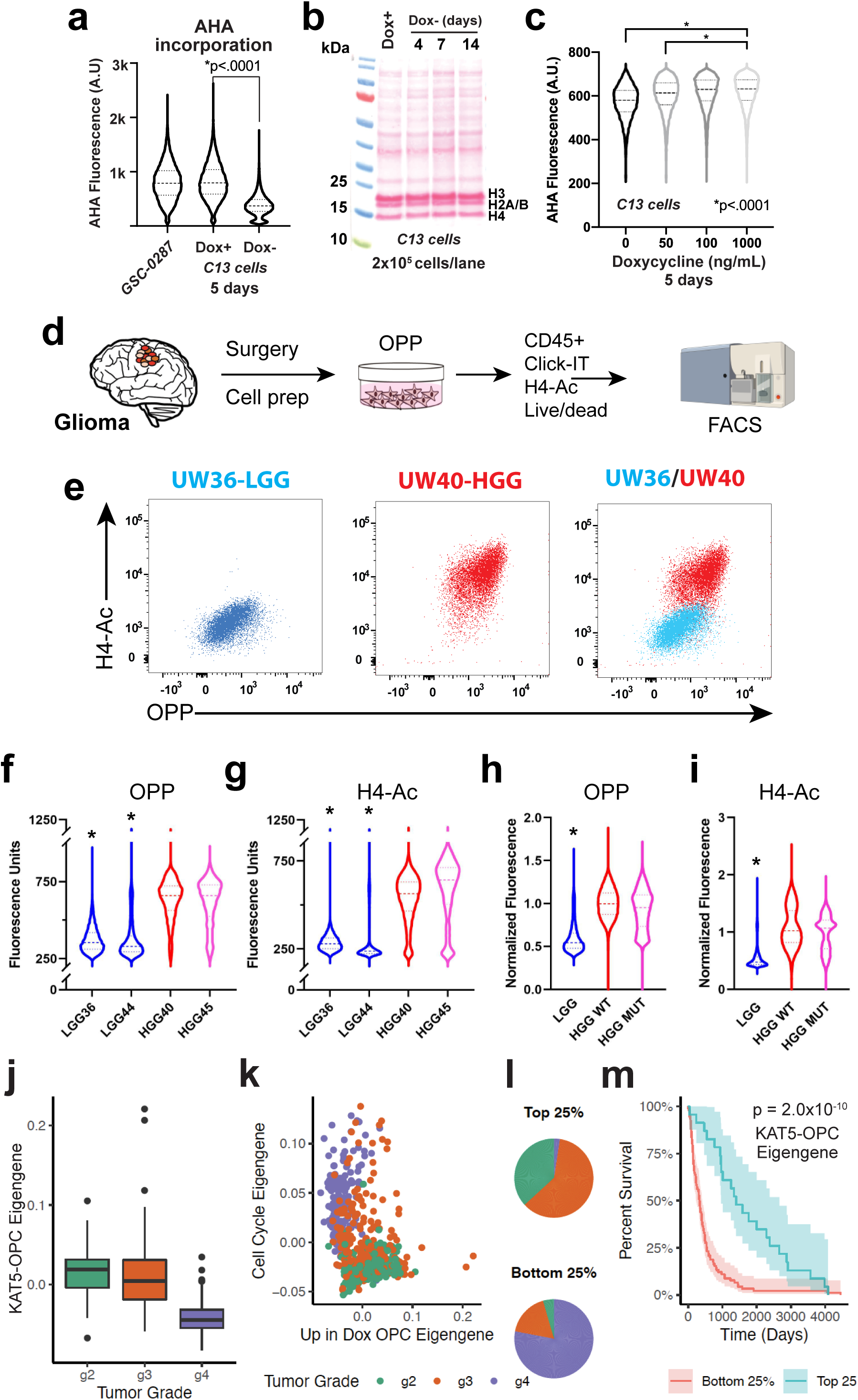
Assessment of KAT5 activity and protein translation rates in primary glioma tumor samples. **a**, Examination of protein synthesis as measured by L-azidohomoalaine (AHA) incorporation for KAT5 inhibited C13 cells after 7 days of Dox withdrawal. KS test was used to test significance. Note: AHA was used for these experiments because OPP (below) cannot be used in puromycin resistant cells. **b**, Examination of total protein content in C13 cells using ponceau staining of total protein extract from 200,000 C13 cells Dox withdrawal of 0, 4, 7, and 14 days. **c**, Examination of protein synthesis via AHA incorporation in C13 cells grown in shown concentrations of Dox for 5 days. KS test was used to test significance. *pval <.0001. **d**, The scheme used for examining protein synthesis using O-propargyl-puromycin (OPP) incorporation in primary tumor cells followed by FACS-based assessment of OPP, histone H4 acetylation (i.e., KAT5 activity), CD45+, and viability. Samples were dissociated, OPP-labeled, viably frozen, thawed, and flow analyzed as a cohort. Tumor cells are CD45-. **e**, Plots of OPP versus pan-H4-Ac in LGG (UW36) and HGG (UW40). **f-g**, Violin plots of OPP and pan-H4-Ac assay results, respectively, from two LGG (IDH1/2mut) (blue), one HGG (IDH1/2wt) (red), and one HGG (IDH1/2 mut) (pink). Each flow event = single cell. KS test was used to assess significance (p<0.0001); LGG vs either HGG WT or IDH1/2mut. **h-i**, Violin plots of OPP and pan-H4-Ac assay results for 3 LGG (IDH1/2mut) (blue), 5 HGG (IDH1/2wt) (red), and two HGG (IDH1/2 mut) (pink) tumors. FACS values are normalized in order for to compare across cohort groups. Each flow event = single cell. KS test was used to assess significance (p<0.0001); LGG vs either HGG WT or IDH1/2mut. **Extended Data Fig. 10** shows similar data for other individual tumors and correlations of H4-Ac vs. OPP incorporation for LGG, HGG, and IDH1/2 mut HGG tumors. **Supplementary Table 16** provides descriptions of each tumor sample used. **j-m**, Analysis of genes affected by KAT5 activity in HGG and LGG tumors. **j**, Relative KAT5-OPC eigengene expression between grade II (n = 226), III (n = 244), and IV (n = 150) tumors (TCGA; LGG and GBM). This eigengene represents the common variation across each patient tumor for the OPC genes upregulated after loss of KAT5 activity in tumors, i.e., first principal component corrected for direction if necessary. All pairwise Student’s t-tests had significant P-values (G2 vs. G3: pval = 0.013; G2 vs G4: pval< 2.2 × 10^-16; G3 vs G4: pval< 2.2 × 10^-16). **k**, Comparison of cell cycle and KAT5-OPC eigengene relative expression in each glioma. Each tumor is colored by its grade (green = II, red = III, and purple = IV). **l**, Distribution of tumor grade between tumors with top 25% and bottom 25% of KAT5-OPC eigengene expression. **m**, Kaplan–Meier survival plot of tumors with top 25% (n = 171) and bottom 25% (n = 171) of KAT5-OPC eigengene expression of KAT5-OPC genes. A Fleming–Harrington survival P-value was used to determine significance. Shaded region is the 95% confidence interval for the survival curve.

We next collected 10 tumors (3 LGG, 5 HGG tumors, and 2 HGG IDH1/2 mutant, which represent recurrent LGG gliomas) and performed KAT5 activity and protein synthesis assays (30 min treatment with O-propargyl-puromycin (OPP)) in match cohorts of tumor cells (**Fig. 7d;**). We used forward and side scatter, CD45+, and viability stains to distinguish tumor cells from brain and immune cells.

These studies revealed that, first, KAT5 activity is dynamic in gliomas: KAT5^hi^ and KAT5^low^ populations are available in each tumor (**Fig. 7e**). Second, in general, LGG tumors have lower H4-Ac levels and protein translation rates than HGG gliomas (**Fig. 7f-I**; **Extended Fig. 10c**). Third, when LGGs recur as HGGs KAT5 activity and protein translation rates are concomitantly up regulated (**Fig. 7f-i; Extended Fig. 10d-f**). Lastly, regardless of grade of tumor, KAT5 activity significantly correlates with protein translation rates in tumor cells, such that low KAT5 correspond to lower protein translation rates and vice versa (**Fig. 7f-i; Extended Fig. 10d-f**).

Of note, similar increases in KAT5 activity were observed in partially transformed human NSCs. Ectopic expression of c-MYC or CyclinD1+ CDK4^R24C^+ dominant-negative p53+TERT^60^ triggered an ~2.5- and ~2-fold increases in H4-Ac staining, respectively, in NSCs, while KAT5^off^ C13 cells had H4-Ac return to levels similar to proliferative NSCs (**Extended Fig. 10g-h**).

### Association of KAT5 regulated genes with gene expression in primary gliomas

We lastly asked whether expression of neurodevelopmental genes upregulated after KAT5 inhibition could be observed in other primary gliomas (TCGA bulk tumor RNA-seq) and among other TCGA tumor types (**Fig. 7j-m**). We observed that expression of these genes is significantly associated with lower grade gliomas and inversely associated with cell cycle gene expression (**Fig. 7j-l**). High expressing tumors also showed better survival outcomes (**Fig. 7m**).

## Discussion

The above studies use a GBM tumor model with a mix of neurodevelopmental subpopulations to determine where tumor cells go after exit from mitosis. Using a G0 reporter, we identify populations of GBM cells reentering the cell cycle and cells with longer G0 resident times, which transition to quasi-neurodevelopmental states associated with oligodendrocyte progenitor and radial glial cell characteristics (**Figs. 1-2**). These states are associated with more indolent, lower grade gliomas and also recently described G0-like gene expression profiles for GBM^5, 30^. We identify KAT5 as a key regulator of these transitions (**Fig. 3**). Its activity prevents GBM from prolonged stays in G0-like states and helps maintain proliferative heterogeneity required for aggressive tumor growth, including invasive behavior. Our studies reveal KAT5-associated transcriptional and epigenetic changes in tumors, which indicate that KAT5 activity biases cell state distribution away from OPC and RG-like subpopulations (**Figs. 4-5**).

Studies in primary glioma samples support these findings. KAT5 activity is dynamic in tumors, where KAT5^low^ tumor cells display quiescent properties (**Fig. 6**). Moreover, tumors with overall lower KAT5 activity (i.e., LGG) have higher expression of genes that are upregulated in KAT5^off^ conditions (**Fig. 7**). This suggests that KAT5 activity is increased in higher grade gliomas, where a larger proportion of cells are in proliferative states. An important future direction is to assay how KAT5 activity alters neurodevelopmental state distribution across lower and higher grade glioma samples.

The KAT5/NuA4 functions as a transcriptional co-activator, whose activities are coordinated with multiple transcription factors: AR/ER^24, 61^, E2F proteins^25^, and MYC in certain contexts^26, 62^. In addition, KAT5/NuA4 complex controls epigenetic regulation of rDNA transcription, which, in turn, ultimately controls ribosome biogenesis and protein translation rates^63^. MYC is also known to regulate Pol I activity and ribosome biogenesis ^64–67^. While loss of KAT5 causes down regulation of MYC protein levels (**Fig. 3j**) and rRNA and protein synthesis levels (**Fig. 7a-c**; **Extended Data Fig. 10a-b**), neither MYC knockdown nor RNA Pol I inhibition (with CX-5641) recapitulated KAT5^off^ phenotypes, causing only modest p27 induction (e.g., <1.6-fold compared >2.5-fold for KAT5 loss) (data not shown).

However, from CUT&TAG analysis, H3K27ac marks AP-1 transcription factor binding sites that are specifically lost in KAT5^off^ tumors (**Fig. 5e**). AP-1 was recently implicated as a regulator of mesenchymal states in GBM^55^. In transformed mouse NSCs, loss of Fosl1 activity causes loss of expression of mesenchymal genes and upregulation of neurodevelopmental genes (e.g., OPC, RG/Ac)^55^. Thus, one possibility is that KAT5 regulates AP-1 activity, directly or indirectly, to block OPC-RG/Ac transitions.

While the above studies suggest how GBM tumor cells can transition from the cell cycle to G0-like states, they do not reveal how tumor cells in G0 states transition back to the cell cycle. However, there are small numbers of cells in G0-like populations that express G1/S genes at higher than background levels expression (e.g., CCNE2 and DNA replication factors). This could indicate that G0 cells become primed to transition back to the cell cycle. Future cell tracking studies will be required to address this and other possibilities.

Is KAT5 a reasonable therapeutic target? Multiple small molecule KAT5 inhibitors have been developed which show promise in sensitizing cancer cells to chemotherapeutics or radiation (rev in^68^). However, in our hands, current KAT5 inhibitors trigger non-specific toxicity in GSCs at effective doses observed for other cell types (not shown). However, it seems likely there does exist a therapeutic window for KAT5 inhibition given its increased activity in HGG (**Fig. 7**). However, the longer-term implications of altering tumor heterogeneity by its inhibition and responses to chemoradiation (i.e., standard-of-care) await further investigation.

Taken together, we establish a model system for the study of G0-like states, identify KAT5 as a critical mediator of G0-like states in GBM, provide evidence for existence of such states in complex model systems and patient samples, and ultimately lay the foundation for therapeutic strategies specifically targeting this population of cells.

## Acknowledgments

We thank members of the Holland, Paddison, Patel, Plaisier, and Tsukiyama labs for helpful discussions, Dr. Atsushi Miyawaki for providing reagents, and Pam Lindberg and An Tyrrell for administrative support. This work was supported by the following grants: Interdisciplinary Training in Cancer Fellowship NCI T32CA080416 (P.H.); Fred Hutch pilot award (A.P., P.P.); NCI/NIH (R01CA190957; P30CA15704) (P.P.); (5R21CA232244) (C.P.); and NINDS/NIH (R01NS119650) (A.P., C.P., P.P.) and Burroughs Wellcome Career Award for Medical Scientists (A.P.).

## Author contributions

Project conception and design was carried out by P.P., A.P., A.M., and H.M.F. Experiments and data analysis were performed by A.M., H.M.F., J.B., K.M., P.H., M.L., and W.J. Critical reagents were generated by M.K., P.C., and L.C.; single cell bioinformatical data analysis and statistics were performed by S.A., S.O., K.J., B.A. and C.L.P. with input from A.P., C.P., and P.P.; P.P., A.P., and A.M. wrote the manuscript with input from other authors.

## Methods

***Key Reagents and Resources are available in Supplementary Table 17.***

### Cell Culture

Patient tumor-derived GSCs were provided by Drs. Jeongwu Lee (Cleveland Clinic), Do-Hyun Nam (Samsung Medical Center, Seoul, Korea), and Steven M. Pollard (University of Edinburgh). Isolates were cultured in NeuroCult NS-A basal medium (StemCell Technologies) supplemented with B27 (Thermo Fisher Scientific), N2 (homemade 2x stock in Advanced DMEM/F-12 (Thermo Fisher Scientific)), EGF and FGF-2 (20 ng/ml) (PeproTech), glutamax (Thermo Fisher Scientific), and antibiotic-antimycotic (Thermo Fisher Scientific). Cells were cultured on laminin (Trevigen or in-house-purified)-coated polystyrene plates and passaged as previously described ^69^, using Accutase (EMD Millipore) to detach cells.

### GSC Tumors

NSG mice (Jackson Labs #005557) used in this study were kept in a 12h light/dark cycle, with food and water ad libitum, in the Fred Hutchinson Cancer Center (FHCC) vivarium. All animal experimental procedures were performed with approval of the FHCC Institutional Animal Care and Use Committee. All procedures followed guidelines outlined in the National Research Council Guide for the Care and Use of Laboratory Animals. 100,000 GSCs were orthotopically xenografted into a single frontal cerebral hemisphere. GSCs were injected using stereo-tactic coordinates: 2 mm lateral from Bregma and 3.5 mm depth and grown for 3-12 weeks according to our previously published protocols ^60, 70, 71^. *<u>Doxycycline dosage of mice</u>:* 827-C13 cells were transplanted into NSG mice that were pre-dosed with Dox (2mg/ml) into the drinking water (supplemented with 5% sucrose) 24 hours in advance. Tumors were allowed to form in the continuous presence of Dox until they were detected by MRI, followed by enrolment into experimental cohorts (i.e., Dox+, Dox-). *<u>EdU pulsing of mice:</u>* 6-20 hrs prior to tumor harvesting mice were intra-peritoneally injected with EdU (100 mg/Kg). *<u>MRI:</u>* volume of interest was manually contoured using T2-weighted brain MRI scans (Bruker 1T scanner) for all animals using Horos Dicom Viewer software (horosproject.org).

### p27 and FUCCI Reporters

The p27 reporter was constructed after (Oki et al., 2014), using a p27 allele that harbors two amino acid substitutions (F62A and F64A) that block binding to Cyclin/CDK complexes but do not interfere with its cell cycle-dependent proteolysis. This p27K^-^ allele was fused to mVenus to create p27K^-^-mVenus. To this end, the p27 allele and mVenus were synthesized as gBlocks (IDT) and cloned via Gibson assembly (NEB) into a modified pGIPz lentiviral expression vector (Open Biosystems). Lentivirally transduced cells were puromycin selected. P27-mVenus reporter cells were sorted for the presence of mVenus on an FACSAria II (BD) and normal growth was verified post-sorting. FUCCI constructs (RIKEN, gift from Dr. Atsushi Miyawaki) were transduced into GSC-0827 cells and sorted sequentially for the presence of mCherry-CDT1(aa30-120) and S/G2/M mAG-Geminin(aa1-110) on an FACSAria II (BD). Normal growth was verified post-sorting and then the FUCCI GSCs were transduced with individual sgRNA-Cas9 and selected with 1 μg/mL puromycin. Cells were grown out for 21 days with splitting every 3-4 days and maintaining equivalent densities. Cells were counted (Nucleocounter NC-100; Eppendorf) and plated 3 days before analysis on an LSR II (BD)

### Lentiviral Production

For virus production, lentiCRISPR v2 plasmids^72^ were transfected using polyethylenimine (Polysciences) into 293T cells along with psPAX and pMD2.G packaging plasmids (Addgene) to produce lentivirus. For the whole-genome CRISPR-Cas9 libraries, 25×150mm plates of 293T cells were seeded at ~15 million cells per plate. Fresh media was added 24 hours later and viral supernatant harvested 24 and 48 hours after that. For screening, virus was concentrated 1000x following ultracentrifugation at 6800*xg* for 20 hours. For validation, lentivirus was used unconcentrated at an MOI<1.

### CRISPR-Cas9 Screening

For large-scale transduction, GSC cells were plated into T225 flasks at an appropriate density such that each replicate had 250-500-fold representation, using the Brunello CRISPR-Cas9 library^73, 74^ (Addgene) (as we have previously published^75^). GSCs were infected at MOI <1 for all cell lines. Cells were infected for 48 hours followed by selection with 2 μg/mL of puromycin for 3 days. Post-selection, a portion of cells were harvested as Day 0 time point. The remaining cells were then passaged in T225 flasks maintaining 250-500-fold representation and cultured for an additional 8 days. Genomic DNA was extracted using QiaAmp Blood Purification Mini or Midi kit (Qiagen). A two-step PCR procedure was performed to amplify sgRNA sequence. For the first PCR, DNA was extracted from the number of cells equivalent to 250-500-fold representation (screen-dependent) for each replicate and the entire sample was amplified for the guide region. For each sample, ~100 separate PCR reactions (library and representation dependent) were performed with 1 μg genomic DNA in each reaction using Herculase II Fusion DNA Polymerase (Agilent) or Phusion High-Fidelity DNA Polymerase (Thermo Fisher). Afterwards, a set of second PCRs was performed to add on Illumina adaptors and to barcode samples, using 10-20ul of the product from the first PCR. Primer sequences are in Supplementary Table 17. We used a primer set to include both a variable 1-6 bp sequence to increase library complexity and 6 bp Illumina barcodes for multiplexing of different biological samples. The whole amplification was carried out with 12 cycles for the first PCR and 18 cycles for the second PCR to maintain linear amplification. Resulting amplicons from the second PCR were column purified using Monarch PCR & DNA Cleanup Kit (New England Biolabs; NEB) to remove genomic DNA and first round PCR product. Purified products were quantified (Qubit 2.0 Fluorometer; Fisher), mixed, and sequenced using HiSeq 2500 (Illumina). Bowtie was used to align the sequenced reads to the guides^76^. The R/Bioconductor package edgeR was used to assess changes across various groups^77^.

### Cas9:sgRNA RNP Nucleofection

Knockout of endogenous genes in GSCs was performed and analyzed as detailed in ^78^. Lyophilized chemically synthesized sgRNA (Synthego) was reconstituted to 100 pmoles/μL in nuclease-free 1X TE Buffer (Tris-EDTA, pH 8.0) and was used directly for RNP complexing or diluted to 30 pmoles/μL in nuclease-free water immediately before use, depending on the particular dosing. Purified sNLS-SpCas9-sNLS (Aldevron) was diluted from 61 pmoles/μL to 10 pmoles/μL in PBS (pH 7.4) immediately before use. To prepare RNP complexes, reconstituted sgRNA was added to SG Cell Line Nucleofector Solution (Lonza), followed by addition of Cas9, to a final volume of 20 μL. A Cas9:sgRNA ratio of 1:2 was used, unless otherwise noted. Total dose of RNPs described in this paper refers to the amount of the limiting complex member (Cas9). The mixture was incubated at room temperature for 15 minutes to allow RNP complexes to form and then placed on ice until use. To nucleofect, 1.3-1.5 x 10^5^ cells were harvested, washed with PBS, resuspended in 20 μL of RNPs, and electroporated using the Amaxa 96-well Shuttle System (Lonza) and program EN-138, similar to^79^. After nucleofection, cells were recovered in pre-warmed culture media and plated onto 12-well or 6-well plates. Media was changed 12-24 hours after nucleofection.

### CRISPR Editing Analysis

Nucleofected cells were harvested at designated timepoints and genomic DNA was extracted (MicroElute Genomic DNA Kit, Omega Bio-Tek). Genomic regions around CRISPR target sites were PCR amplified using primers located (whenever possible) at least 250bp outside cut sites. After size verification by agarose gel electrophoresis, PCR products were column-purified (Monarch PCR & DNA Clean-up Kit, New England BioLabs) and submitted for Sanger sequencing (Genewiz) using unique sequencing primers. The resulting trace files for edited cells versus control cells (nucleofected with non-targeting Cas9:sgRNA) were analyzed for predicted indel composition using the ICE web tool ^80^.

### Flow Cytometry

GSC cells that incorporated EdU while alive (2-24 hrs, 10-2 μM) EU (1μM for 1hr) or AHA (100 μM for 1 hr after cells were grown in Methionine-free SILAC media for 30 min.) were fixed (4% PFA) and permeabilized (0.1% Saponin) and subjected to Click-iT Plus chemistry detection prior to analysis by flow cytometry. In some cases, the same cells were also stained for H4-panAc (1:100) for KAT5 function evaluation, DAPI (0.001 μg/ul) for DNA content analysis, or Pyronin Y for RNA content analysis. H4-panAc staining was done in the presence of 0.3% Triton X-100 and 5% normal goat serum for 1 hr at room temperature and a secondary antibody conjugated to an Alexa Fluor was use for primary antibody detection (for 30 min. at room temperature) during flow analysis. Processed cells were flow cytometry analyzed immediately using either a BD FACSymphony A5 or BD LSRFortessa X-50 machine. Results were analyzed using FlowJo software. For experiments using sorted p27-high populations from GSC-p27-mVenus reporter cells, a gate at ~20% p27 high was used to isolate these populations. Either a BD FACSymphony S6 or Sony MA900 cell sorters were used.

### Histone H4 Acetylation and OPP Analysis of Primary Glioma Samples

Over the course of 2 years, 10 patient glioma tumors were collected and assayed: 3 LGGs (UW33, UW36, UW44), 5 HGGs (UW27, UW31, UW34, UW38, UW40), and 2 HGGs IDH1/2 mutant (UW26, UW44), which represent recurrent LGGs (**Supplementary Table 16**). Freshly resected tumors were dissociated by a combination of mechanical and enzymatic methods using a brain tumor dissociation kit (Miltenyi # 130-095-942) according to the manufacture’s protocol. Cells were counted, and while alive, they were incubated with OPP (2 μM final, 1 million cells in 1 ml NSC media, in a low binding tube at 37°C for 30 min). Then cells were washed in warm NSC, and slowly frozen (1°C cooling/1 min.) in NSC media supplemented with 10% DMSO. Cohorts of HGG^WT^ and LGG and/or HGG^MUT^ were processed together in order to use the HGG^WT^ as normalization among cohorts that were collected over 2 years. Frozen cells were thawed using a method that recovers a large percentage of viable cells after freeze/thaw (frozen cells were thawed at 37°C for 2 min., then warm NSC media was added dropwise to cells by doubling the volume every minute to a total volume of 32 ml, starting from one 1ml frozen cell vial). While cells were alive, a CD45 antibody was used to surface stain immune tumor populations, a viability fixable Zombie dye (BioLegend) was used to determine live cells, then fixed in 4% PFA, permeabilized, and processed for Click-iT chemistry to detect OPP, and intracellularly stained for H4Ac. Processed cells were flow cytometry analyzed immediately using either a BD FACSymphony A5 or BD LSRFortessa X-50 machine. FSC-A and SSC-A plots were used to gate the tumor populations and eliminate cell debris, FSC-A and FSC-H plots were used to gate singlet populations, a gate on Zombie dye negative cells was used to isolate live cells, followed by a CD45 negative gate to isolate the non-immune tumor cell populations that were entered into the downstream analysis to evaluate protein synthesis rates (OPP+), and KAT5 activity (H4Ac+).

### Western blotting

Cells were harvested, washed with PBS, and either immediately lysed or snap-frozen and stored at −80°C until lysis. Cells were lysed with RIPA buffer (Thermo Scientific cat# 89900), 1X complete protease inhibitor cocktail (complete Mini EDTA-free, Roche) and 2.5U/μL benzonase nuclease (Novagen) in RIPA buffer supplemented with 1 mM final MgCl_2_ concentration, at room temperature for 15 minutes. To enhance histone modification detection, a total histone extraction kit was used according to the manufacturer’s protocol (Epigentek # OP-0006). Cell lysates were quantified using Pierce BCA protein assay reagent and proteins were loaded onto SDS-PAGE for western blot. The Trans-Blot Turbo transfer system (Bio-Rad) was used according to the manufacturer’s instructions. Ponceau S was used to visualize total proteins on the western blot membranes prior to the blocking step. An Odyssey infrared imaging system was used to visualize blots (LI-COR) following the manufacturer’s instructions.

### Creation of Doxycycline controllable KAT5 GSC-0827 cells

A KAT5-V5 tag ORF was cloned into the Tet-inducible expression retroviral plasmid pTight-TURB vector (MSCV-tetO7-mCherry-UBC-rtTA-Blast) via Gibson assembly a.GSC-0827/p27-mVenus reporter cells were infected with pTight-TURB-KAT5-V5 (3 rounds of infection over a 3 day period) and Blasticidin selected. To turn on the Tet-inducible expression, cells were grown in 1 μg/ml Doxycycline (Dox) and nucleofected with sgKAT5:Cas9 complexes to KO endogenous KAT5 (using an sgRNA that spans an exon-intron junction that does not recognize the KAT5 ORF). After a 72hr recovery period, cells plated in 96 well plates at a frequency of ~.25 cells per well in the presence of Dox and allowed to outgrow for two weeks with media changes every three day. Afterwards, several clones that preserved the p27-mVenus reporter were picked and evaluated for growth arrest and p27 reporter induction upon Dox withdrawal. Clone 13, used above, displayed the most uniform Dox+ growth and Dox-growth arrest among ~10 clones evaluated.

### GSC invasion assays

GSC-0827 C13 cells were cultured with 1 μg/ml Doxycycline or without Doxycycline for 4 days or 7 days. A total of 5X10^3^ cells were seeded in an ultra-low attachment round bottom 96-well plate (Corning, 7007) and incubated 3 days. For the invasion assay, an 8-well chamber slide (LAB-TEK, 154534) was coated with 30 μg/ml Matrigel (Corning, 356231) and incubated at 4 °C overnight, followed by washing with ice-cold PBS. Then, 3-5 spheroids were transferred into 150 μl of 7.8 mg/ml Matrigel-covered wells and incubated at 37 °C for 1 hour. After polymerization, 150 μl NSC media were overlaid with or without 2 μg/ml Doxycycline as needed. Phase-contrast images were captured every 24 hours for 72 hours on a Nikon Eclipse TS100 microscope. The area covered by invading cells was measured using FIJI.

The average cell invasion distance for each spheroid was calculated using the following formula: Average cell migration 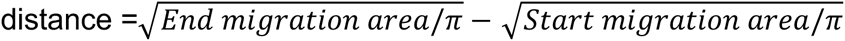.

### scRNA-seq analysis

Single cell RNA-sequencing was performed using 10x Genomics’ reagents, instruments, and protocols. Single cell RNA-Seq libraries were prepared using Chromium Single Cell 3ʹ Reagent Kit. CellRanger^81^(v5.0 from 10× Genomics) was used to align, quantify, and provide basic quality control metrics for the scRNA-seq data. Souporcell^82^ was used to deconvolute scRNA-seq data for each GSC cell line. Using Seurat^83^ (version 4), the scRNA-seq data was normalized using the SCTransform pipeline and were merged the GSC 827 tumor replicates to build an integrated reference. FindTransferAnchors and MapQuery from Seurat, was used to map the query tumors to the integrated reference. FindAllMarkers was used to find differentially expressed genes for each cluster of each tumor. AddModuleScore from Seurat to calculate the average expression levels of different gene lists of interest for each tumor type. ggplot2 was visualize to make bar plots to visualize the number of genes and cells in each cluster. ccSeurat^83^ and ccAF^5^ were used to score cell cycle states for each cell. scVelo^11^ was used to perform velocity analysis. The ToppCell Atlas ^10^ was used to perform gene set enrichment analysis on each of the differentially expressed gene list from each cluster.

### CUT&Tag analysis

CUT&Tag assays were performed using the EpiCypher CUTANA CUT&Tag kit (Cat# 14-1101) according to manufacturer’s instructions. We used 100,000 cells as input per each sample and proceeded with nuclei extraction before binding to ConA beads. The samples were derived from GSC-0827 C13 orthotopic xenografts that were also used for scRNA-seq analysis in this study. CUT&Tag analysis was assayed for the following epigenetic markers: H3K4Me2 (Millipore, Cat# 07-030), H3K27Ac (Millipore, Cat# MABE647) and H3K27Me3 (Cell Signaling Technologies, Cat# 9733).

The single-cell RNA sequencing data files are available on the GEO database at GSE198524. [Review token access : wtehwmkonvypbyv].

The code used to process and analyze the data is available at https://github.com/sonali bioc/GSC_scRNASeq_KAT5paper. All other data associated with this study are present in supplementary materials and tables.

## References

1 Patel, A. P. et al. Single-cell RNA-seq highlights intratumoral heterogeneity in primary glioblastoma. Science 344, 1396–1401 (2014). https://doi.org:10.1126/science.1254257

2 Neftel, C. et al. An Integrative Model of Cellular States, Plasticity, and Genetics for Glioblastoma. Cell 178, 835–849 e821 (2019). https://doi.org:10.1016/j.cell.2019.06.024

3 Hanahan, D. Hallmarks of Cancer: New Dimensions. Cancer discovery 12, 31–46 (2022). https://doi.org:10.1158/2159-8290.CD-21-1059

4 Lathia, J. D., Mack, S. C., Mulkearns-Hubert, E. E., Valentim, C. L. & Rich, J. N. Cancer stem cells in glioblastoma. Genes Dev 29, 1203–1217 (2015). https://doi.org:10.1101/gad.261982.115

5 O’Connor, S. A. et al. Neural G0: a quiescent-like state found in neuroepithelial-derived cells and glioma. Molecular systems biology 17, e9522 (2021). https://doi.org:10.15252/msb.20209522

6 Llorens-Bobadilla, E. et al. Single-Cell Transcriptomics Reveals a Population of Dormant Neural Stem Cells that Become Activated upon Brain Injury. Cell Stem Cell 17, 329–340 (2015). https://doi.org:10.1016/j.stem.2015.07.002

7 Artegiani, B. et al. A Single-Cell RNA Sequencing Study Reveals Cellular and Molecular Dynamics of the Hippocampal Neurogenic Niche. Cell Rep 21, 3271–3284 (2017). https://doi.org:10.1016/j.celrep.2017.11.050

8 Dahlrot, R. H. et al. Prognostic role of Ki-67 in glioblastomas excluding contribution from non-neoplastic cells. Scientific reports 11, 17918 (2021). https://doi.org:10.1038/s41598-021-95958-9

9 Becht, E. et al. Dimensionality reduction for visualizing single-cell data using UMAP. Nat Biotechnol (2018). https://doi.org:10.1038/nbt.4314

10 Jin, K. et al. An Interactive Single Cell Web Portal Identifies Gene and Cell Networks in COVID-19 Host Responses. iScience, 103115 (2021). https://doi.org:10.1016/j.isci.2021.103115

11 Bergen, V., Lange, M., Peidli, S., Wolf, F. A. & Theis, F. J. Generalizing RNA velocity to transient cell states through dynamical modeling. Nat Biotechnol 38, 1408–1414 (2020). https://doi.org:10.1038/s41587-020-0591-3

12 Fischer, M., Schade, A. E., Branigan, T. B., Muller, G. A. & DeCaprio, J. A. Coordinating gene expression during the cell cycle. Trends in biochemical sciences 47, 1009–1022 (2022). https://doi.org:10.1016/j.tibs.2022.06.007

13 Darzynkiewicz, Z., Traganos, F. & Melamed, M. R. New cell cycle compartments identified by multiparameter flow cytometry. Cytometry 1, 98–108 (1980). https://doi.org:10.1002/cyto.990010203

14 McKnight, J. N., Boerma, J. W., Breeden, L. L. & Tsukiyama, T. Global Promoter Targeting of a Conserved Lysine Deacetylase for Transcriptional Shutoff during Quiescence Entry. Mol Cell 59, 732–743 (2015). https://doi.org:10.1016/j.molcel.2015.07.014

15 Park, J. S. et al. N-myc downstream regulated gene 1 (ndrg1) functions as a molecular switch for cellular adaptation to hypoxia. eLife 11 (2022). https://doi.org:10.7554/eLife.74031

16 Levy, A. P., Levy, N. S., Wegner, S. & Goldberg, M. A. Transcriptional regulation of the rat vascular endothelial growth factor gene by hypoxia. J Biol Chem 270, 13333–13340 (1995). https://doi.org:10.1074/jbc.270.22.13333

17 Spencer, S. L. et al. The proliferation-quiescence decision is controlled by a bifurcation in CDK2 activity at mitotic exit. Cell 155, 369–383 (2013). https://doi.org:10.1016/j.cell.2013.08.062

18 Boele, J. et al. PAPD5-mediated 3’ adenylation and subsequent degradation of miR-21 is disrupted in proliferative disease. Proc Natl Acad Sci U S A 111, 11467–11472 (2014). https://doi.org:10.1073/pnas.1317751111

19 Fok, W. C. et al. Posttranscriptional modulation of TERC by PAPD5 inhibition rescues hematopoietic development in dyskeratosis congenita. Blood 133, 1308–1312 (2019). https://doi.org:10.1182/blood-2018-11-885368

20 Sinturel, F. et al. Diurnal Oscillations in Liver Mass and Cell Size Accompany Ribosome Assembly Cycles. Cell 169, 651–663 e614 (2017). https://doi.org:10.1016/j.cell.2017.04.015

21 Johnston, G. C., Pringle, J. R. & Hartwell, L. H. Coordination of growth with cell division in the yeast Saccharomyces cerevisiae. Exp Cell Res 105, 79–98 (1977). https://doi.org:10.1016/0014-4827(77)90154-9

22 Jorgensen, P. & Tyers, M. How cells coordinate growth and division. Curr Biol 14, R1014–1027 (2004). https://doi.org:10.1016/j.cub.2004.11.027

23 Dulken, B. W., Leeman, D. S., Boutet, S. C., Hebestreit, K. & Brunet, A. Single-Cell Transcriptomic Analysis Defines Heterogeneity and Transcriptional Dynamics in the Adult Neural Stem Cell Lineage. Cell Rep 18, 777–790 (2017). https://doi.org:10.1016/j.celrep.2016.12.060

24 Jeong, K. W. et al. Recognition of enhancer element-specific histone methylation by TIP60 in transcriptional activation. Nat Struct Mol Biol 18, 1358–1365 (2011). https://doi.org:10.1038/nsmb.2153

25 Taubert, S. et al. E2F-dependent histone acetylation and recruitment of the Tip60 acetyltransferase complex to chromatin in late G1. Mol Cell Biol 24, 4546–4556 (2004). https://doi.org:10.1128/MCB.24.10.4546-4556.2004

26 Ravens, S., Yu, C., Ye, T., Stierle, M. & Tora, L. Tip60 complex binds to active Pol II promoters and a subset of enhancers and co-regulates the c-Myc network in mouse embryonic stem cells. Epigenetics & chromatin 8, 45 (2015). https://doi.org:10.1186/s13072-015-0039-z

27 Xu, Y. et al. The p400 ATPase regulates nucleosome stability and chromatin ubiquitination during DNA repair. J Cell Biol 191, 31–43 (2010). https://doi.org:10.1083/jcb.201001160

28 Courilleau, C. et al. The chromatin remodeler p400 ATPase facilitates Rad51-mediated repair of DNA double-strand breaks. J Cell Biol 199, 1067–1081 (2012). https://doi.org:10.1083/jcb.201205059

29 Osuka, S. et al. N-cadherin upregulation mediates adaptive radioresistance in glioblastoma. J Clin Invest 131 (2021). https://doi.org:10.1172/JCI136098

30 Xie, X. P. et al. Quiescent human glioblastoma cancer stem cells drive tumor initiation, expansion, and recurrence following chemotherapy. Dev Cell 57, 32–46 e38 (2022). https://doi.org:10.1016/j.devcel.2021.12.007

31 Zhang, L. et al. Pleiotrophin enhances PDGFB-induced gliomagenesis through increased proliferation of neural progenitor cells. Oncotarget 7, 80382–80390 (2016). https://doi.org:10.18632/oncotarget.12983

32 Fujikawa, A. et al. Small-molecule inhibition of PTPRZ reduces tumor growth in a rat model of glioblastoma. Scientific reports 6, 20473 (2016). https://doi.org:10.1038/srep20473

33 Fujikawa, A. et al. Targeting PTPRZ inhibits stem cell-like properties and tumorigenicity in glioblastoma cells. Scientific reports 7, 5609 (2017). https://doi.org:10.1038/s41598-017-05931-8

34 Knudsen, A. M. et al. Surgical resection of glioblastomas induces pleiotrophin-mediated self-renewal of glioblastoma stem cells in recurrent tumors. Neuro Oncol 24, 1074–1087 (2022). https://doi.org:10.1093/neuonc/noab302

35 Brozzi, F., Arcuri, C., Giambanco, I. & Donato, R. S100B Protein Regulates Astrocyte Shape and Migration via Interaction with Src Kinase: IMPLICATIONS FOR ASTROCYTE DEVELOPMENT, ACTIVATION, AND TUMOR GROWTH. J Biol Chem 284, 8797–8811 (2009). https://doi.org:10.1074/jbc.M805897200

36 Wang, H. et al. S100B promotes glioma growth through chemoattraction of myeloid-derived macrophages. Clin Cancer Res 19, 3764–3775 (2013). https://doi.org:10.1158/1078-0432.CCR-12-3725

37 Golembieski, W. A. et al. HSP27 mediates SPARC-induced changes in glioma morphology, migration, and invasion. Glia 56, 1061–1075 (2008). https://doi.org:10.1002/glia.20679

38 Shi, Q. et al. Targeting SPARC expression decreases glioma cellular survival and invasion associated with reduced activities of FAK and ILK kinases. Oncogene 26, 4084–4094 (2007). https://doi.org:10.1038/sj.onc.1210181

39 Qin, E. Y. et al. Neural Precursor-Derived Pleiotrophin Mediates Subventricular Zone Invasion by Glioma. Cell 170, 845–859 e819 (2017). https://doi.org:10.1016/j.cell.2017.07.016

40 Kim, J. et al. Ttyh1 regulates embryonic neural stem cell properties by enhancing the Notch signaling pathway. EMBO Rep 19 (2018). https://doi.org:10.15252/embr.201745472

41 Wu, H. N. et al. Deficiency of Ttyh1 downstream to Notch signaling results in precocious differentiation of neural stem cells. Biochem Biophys Res Commun 514, 842–847 (2019). https://doi.org:10.1016/j.bbrc.2019.04.181

42 Bhat, K. P. et al. Mesenchymal differentiation mediated by NF-kappaB promotes radiation resistance in glioblastoma. Cancer Cell 24, 331–346 (2013). https://doi.org:10.1016/j.ccr.2013.08.001

43 Kimura, A. & Horikoshi, M. Tip60 acetylates six lysines of a specific class in core histones in vitro. Genes Cells 3, 789–800 (1998).

44 Kaya-Okur, H. S. et al. CUT&Tag for efficient epigenomic profiling of small samples and single cells. Nature communications 10, 1930 (2019). https://doi.org:10.1038/s41467-019-09982-5

45 Creyghton, M. P. et al. Histone H3K27ac separates active from poised enhancers and predicts developmental state. Proc Natl Acad Sci U S A 107, 21931–21936 (2010). https://doi.org:10.1073/pnas.1016071107

46 Boyer, L. A., Mathur, D. & Jaenisch, R. Molecular control of pluripotency. Current Opinion in Genetics & Development 16, 455–462 (2006).

47 Margueron, R. & Reinberg, D. The Polycomb complex PRC2 and its mark in life. Nature 469, 343–349 (2011). https://doi.org:10.1038/nature09784

48 Ernst, J. & Kellis, M. Chromatin-state discovery and genome annotation with ChromHMM. Nat Protoc 12, 2478–2492 (2017). https://doi.org:10.1038/nprot.2017.124

49 Zhang, R. et al. Id4 Downstream of Notch2 Maintains Neural Stem Cell Quiescence in the Adult Hippocampus. Cell Rep 28, 1485–1498 e1486 (2019). https://doi.org:10.1016/j.celrep.2019.07.014

50 Liu, F. et al. EGFR Mutation Promotes Glioblastoma through Epigenome and Transcription Factor Network Remodeling. Mol Cell 60, 307–318 (2015). https://doi.org:10.1016/j.molcel.2015.09.002

51 Augustus, M. et al. Identification of CRYAB(+) KCNN3(+) SOX9(+) Astrocyte-Like and EGFR(+) PDGFRA(+) OLIG1(+) Oligodendrocyte-Like Tumoral Cells in Diffuse IDH1-Mutant Gliomas and Implication of NOTCH1 Signalling in Their Genesis. Cancers (Basel) 13 (2021). https://doi.org:10.3390/cancers13092107

52 Moreira, F. et al. NPAS3 demonstrates features of a tumor suppressive role in driving the progression of Astrocytomas. The American journal of pathology 179, 462–476 (2011). https://doi.org:10.1016/j.ajpath.2011.03.044

53 Hou, L., Srivastava, Y. & Jauch, R. Molecular basis for the genome engagement by Sox proteins. Seminars in cell & developmental biology 63, 2–12 (2017). https://doi.org:10.1016/j.semcdb.2016.08.005

54 Serresi, M. et al. Functional antagonism of chromatin modulators regulates epithelial-mesenchymal transition. Sci Adv 7 (2021). https://doi.org:10.1126/sciadv.abd7974

55 Marques, C. et al. NF1 regulates mesenchymal glioblastoma plasticity and aggressiveness through the AP-1 transcription factor FOSL1. eLife 10 (2021). https://doi.org:10.7554/eLife.64846

56 Whyte, W. A. et al. Master transcription factors and mediator establish super-enhancers at key cell identity genes. Cell 153, 307–319 (2013). https://doi.org:10.1016/j.cell.2013.03.035

57 Eckel-Passow, J. E. et al. Glioma Groups Based on 1p/19q, IDH, and TERT Promoter Mutations in Tumors. N Engl J Med 372, 2499–2508 (2015). https://doi.org:10.1056/NEJMoa1407279

58 Parsons, D. W. et al. An integrated genomic analysis of human glioblastoma multiforme. Science 321, 1807–1812 (2008). https://doi.org:1164382 [pii] 10.1126/science.1164382

59 Ohgaki, H. & Kleihues, P. The definition of primary and secondary glioblastoma. Clin Cancer Res 19, 764–772 (2013). https://doi.org:10.1158/1078-0432.CCR-12-3002

60 Hubert, C. G. et al. Genome-wide RNAi screens in human brain tumor isolates reveal a novel viability requirement for PHF5A. Genes Dev 27, 1032–1045 (2013). https://doi.org:10.1101/gad.212548.112

61 Brady, M. E. et al. Tip60 is a nuclear hormone receptor coactivator. J Biol Chem 274, 17599–17604 (1999). https://doi.org:10.1074/jbc.274.25.17599

62 Kim, J. et al. A Myc network accounts for similarities between embryonic stem and cancer cell transcription programs. Cell 143, 313–324 (2010). https://doi.org:10.1016/j.cell.2010.09.010

63 Halkidou, K., Logan, I. R., Cook, S., Neal, D. E. & Robson, C. N. Putative involvement of the histone acetyltransferase Tip60 in ribosomal gene transcription. Nucleic Acids Res 32, 1654–1665 (2004). https://doi.org:10.1093/nar/gkh296

64 Arabi, A. et al. c-Myc associates with ribosomal DNA and activates RNA polymerase I transcription. Nat Cell Biol 7, 303–310 (2005). https://doi.org:10.1038/ncb1225

65 Dai, M. S. & Lu, H. Crosstalk between c-Myc and ribosome in ribosomal biogenesis and cancer. Journal of cellular biochemistry 105, 670–677 (2008). https://doi.org:10.1002/jcb.21895

66 Grandori, C. et al. c-Myc binds to human ribosomal DNA and stimulates transcription of rRNA genes by RNA polymerase I. Nat Cell Biol 7, 311–318 (2005). https://doi.org:10.1038/ncb1224

67 Shiue, C. N., Berkson, R. G. & Wright, A. P. c-Myc induces changes in higher order rDNA structure on stimulation of quiescent cells. Oncogene 28, 1833–1842 (2009). https://doi.org:10.1038/onc.2009.21

68 Brown, J. A., Bourke, E., Eriksson, L. A. & Kerin, M. J. Targeting cancer using KAT inhibitors to mimic lethal knockouts. Biochem Soc Trans 44, 979–986 (2016). https://doi.org:10.1042/BST20160081

69 Pollard, S. M. et al. Glioma stem cell lines expanded in adherent culture have tumor-specific phenotypes and are suitable for chemical and genetic screens. Cell Stem Cell 4, 568–580 (2009). https://doi.org:10.1016/j.stem.2009.03.014

70 Toledo, C. M. et al. BuGZ is required for Bub3 stability, Bub1 kinetochore function, and chromosome alignment. Dev Cell 28, 282–294 (2014). https://doi.org:10.1016/j.devcel.2013.12.014

71 Ding, Y. et al. Cancer-Specific requirement for BUB1B/BUBR1 in human brain tumor isolates and genetically transformed cells. Cancer discovery 3, 198–211 (2013). https://doi.org:10.1158/2159-8290.CD-12-0353

72 Sanjana, N. E., Shalem, O. & Zhang, F. Improved vectors and genome-wide libraries for CRISPR screening. Nat Methods 11, 783–784 (2014). https://doi.org:10.1038/nmeth.3047

73 Shalem, O. et al. Genome-scale CRISPR-Cas9 knockout screening in human cells. Science 343, 84–87 (2014). https://doi.org:10.1126/science.1247005

74 Doench, J. G. et al. Optimized sgRNA design to maximize activity and minimize off-target effects of CRISPR-Cas9. Nat Biotechnol 34, 184–191 (2016). https://doi.org:10.1038/nbt.3437

75 Toledo, C. M. et al. Genome-wide CRISPR-Cas9 Screens Reveal Loss of Redundancy between PKMYT1 and WEE1 in Glioblastoma Stem-like Cells. Cell Rep 13, 2425–2439 (2015). https://doi.org:10.1016/j.celrep.2015.11.021

76 Langmead, B., Trapnell, C., Pop, M. & Salzberg, S. L. Ultrafast and memory-efficient alignment of short DNA sequences to the human genome. Genome Biol 10, R25 (2009). https://doi.org:gb-2009-10-3-r25 [pii] 10.1186/gb-2009-10-3-r25

77 Robinson, M. D., McCarthy, D. J. & Smyth, G. K. edgeR: a Bioconductor package for differential expression analysis of digital gene expression data. Bioinformatics 26, 139–140 (2010). https://doi.org:btp616 [pii] 10.1093/bioinformatics/btp616

78 Hoellerbauer, P., Kufeld, M. & Paddison, P. J. Efficient Multi-Allelic Genome Editing of Primary Cell Cultures via CRISPR-Cas9 Ribonucleoprotein Nucleofection. Current protocols in stem cell biology 54, e126 (2020). https://doi.org:10.1002/cpsc.126

79 Bressan, R. B. et al. Efficient CRISPR/Cas9-assisted gene targeting enables rapid and precise genetic manipulation of mammalian neural stem cells. Development 144, 635–648 (2017). https://doi.org:10.1242/dev.140855

80 Hsiau, T., et al. Inference of CRISPR Edits from Sanger Trace Data. bioRxiv (2019).

81 Zheng, G. X. et al. Massively parallel digital transcriptional profiling of single cells. Nature communications 8, 14049 (2017). https://doi.org:10.1038/ncomms14049

82 Heaton, H. et al. Souporcell: robust clustering of single-cell RNA-seq data by genotype without reference genotypes. Nat Methods 17, 615–620 (2020). https://doi.org:10.1038/s41592-020-0820-1

83 Butler, A., Hoffman, P., Smibert, P., Papalexi, E. & Satija, R. Integrating single-cell transcriptomic data across different conditions, technologies, and species. Nat Biotechnol 36, 411–420 (2018). https://doi.org:10.1038/nbt.4096

84 Ding, Y. et al. ZNF131 suppresses centrosome fragmentation in glioblastoma stem-like cells through regulation of HAUS5. Oncotarget (2017). https://doi.org:10.18632/oncotarget.18153

85 Satija, R., Farrell, J. A., Gennert, D., Schier, A. F. & Regev, A. Spatial reconstruction of single-cell gene expression data. Nat Biotechnol 33, 495–502 (2015). https://doi.org:10.1038/nbt.3192

86 Arora, M., Moser, J., Phadke, H., Basha, A. A. & Spencer, S. L. Endogenous Replication Stress in Mother Cells Leads to Quiescence of Daughter Cells. Cell Rep 19, 1351–1364 (2017). https://doi.org:10.1016/j.celrep.2017.04.055

