## Extended Data and Supplementary Figures for "KAT5 regulates neurodevelopmental states associated with G0-like populations in glioblastoma"

### Extended Data Figure 1

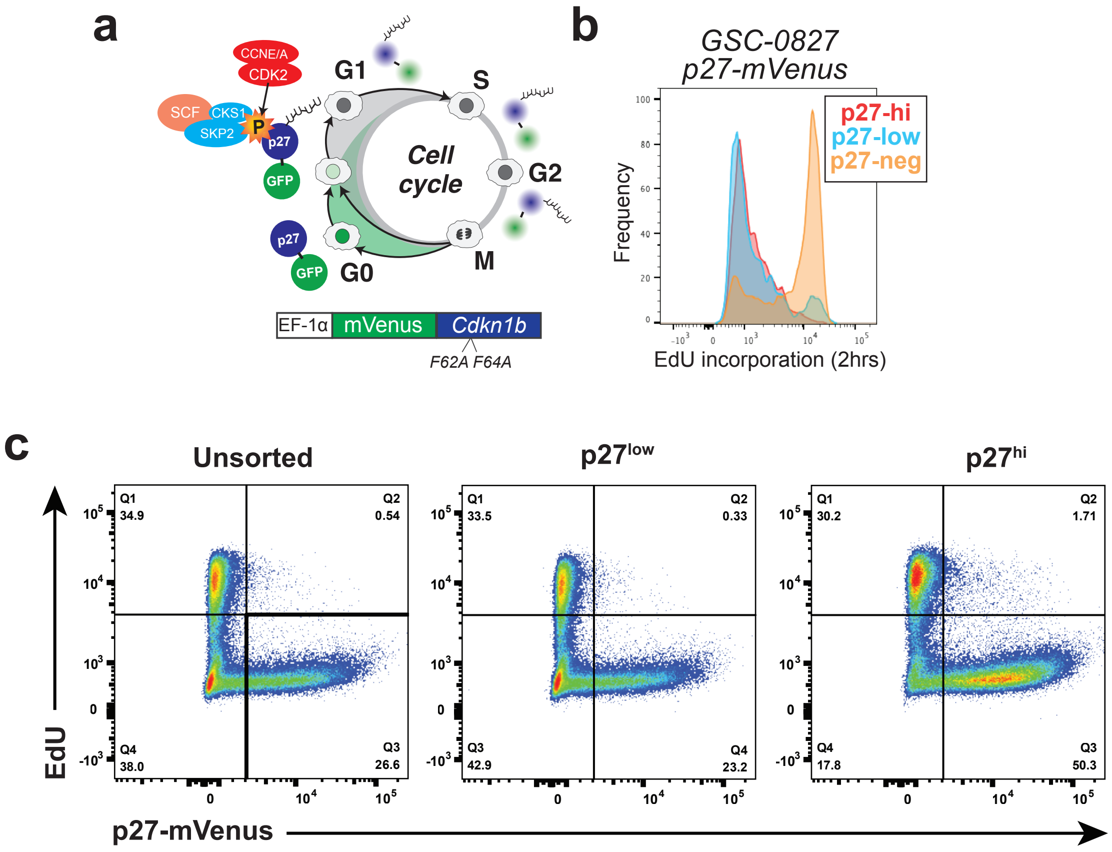

#### Extended Data Figure Legends

**Extended Data Figure 1**: Validation of G0-reporter in GSC-0827 cells.
**a**, The p27-mVenus G0/quiescence reporter and its regulation during the cell cycle. For S/G2/M phases of the cell cycle, p27 is targeted for proteolysis by the $SCF^{Skp2}$ E3 ubiquitin ligase complex (Chu et al., 2008). p27 is additionally regulated in G1 via targeted degradation by the Kip1 ubiquitylation-promoting complex at the G0-G1 transition (Chu et al., 2008). As a result, p27 protein only accumulates during G0. The p27 reporter was constructed with a p27 allele harboring two amino acid substitutions (F62A and F64A) that block binding to Cyclin/CDK complexes (preventing functional activity) but do not interfere with its cell cycle-dependent proteolysis (Oki et al., 2014).
**b**, Flow-based examination of EdU incorporation (2hrs) of p27 high, low, and negative GSC-0827 cells using the p27-mVenus reporter. Consistent with reporting of G0-like states in GSCs, EdU incorporation is significantly suppressed in p27-mVenus+ and especially p27-mVenushi cells. p27-mVenushi cells= top 20%. p27-mVenuslow cells = bottom 20%.
**c**, DNA replication and p27 assay for outgrown populations of p27hi sorted GSC-0827s. Controls include p27low sorted and non-sorted cells. To ensure that p27hi cells could re-enter the cell cycle, p27hi cells were sorted, recultured, and assayed 7 days later for EdU incorporation and p27-mVenus levels. The p27hi cells showed high, but somewhat diminished, EdU incorporation rate of 30% versus 35% for control cells, and also had higher residual p27-mVenus expression. These results are consistent with a preponderance of p27hi cells being division capable and entering the cell cycle with delayed and somewhat variable kinetics as cells exit from G0.

### Extended Data Figure 2

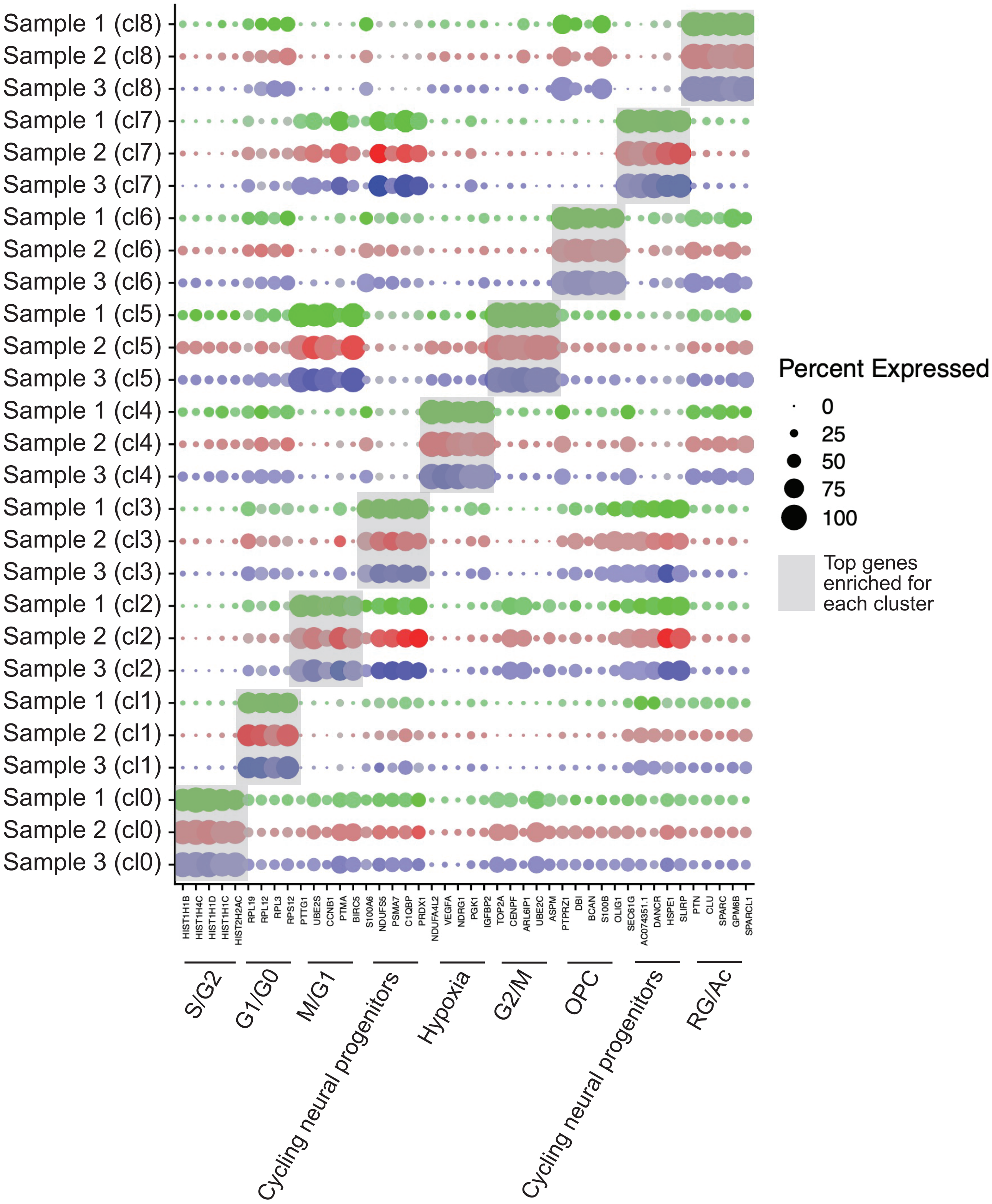

**Extended Data Figure 2**: Dot plot top differentially expressed genes in UMAP clusters for GSC-0827 tumor scRNA-seq data, associated with **Figure 1**. The data is derived from 3 scRNA-seq data sets from GSC-0827 tumors (i.e., samples 1-3). The samples are associated by cluster (cl) from cl0 to cl8. Genes were ranked based on adjusted p-values from differentiation expression analysis of for each cluster **Supplementary Table 1**). The size of the dots corresponds to the proportion of cells within the cluster expressing the gene (% Exp.). Cluster designations from **Figure 1b** are shown below each cluster-specific gene set.

### Extended Data Figure 3

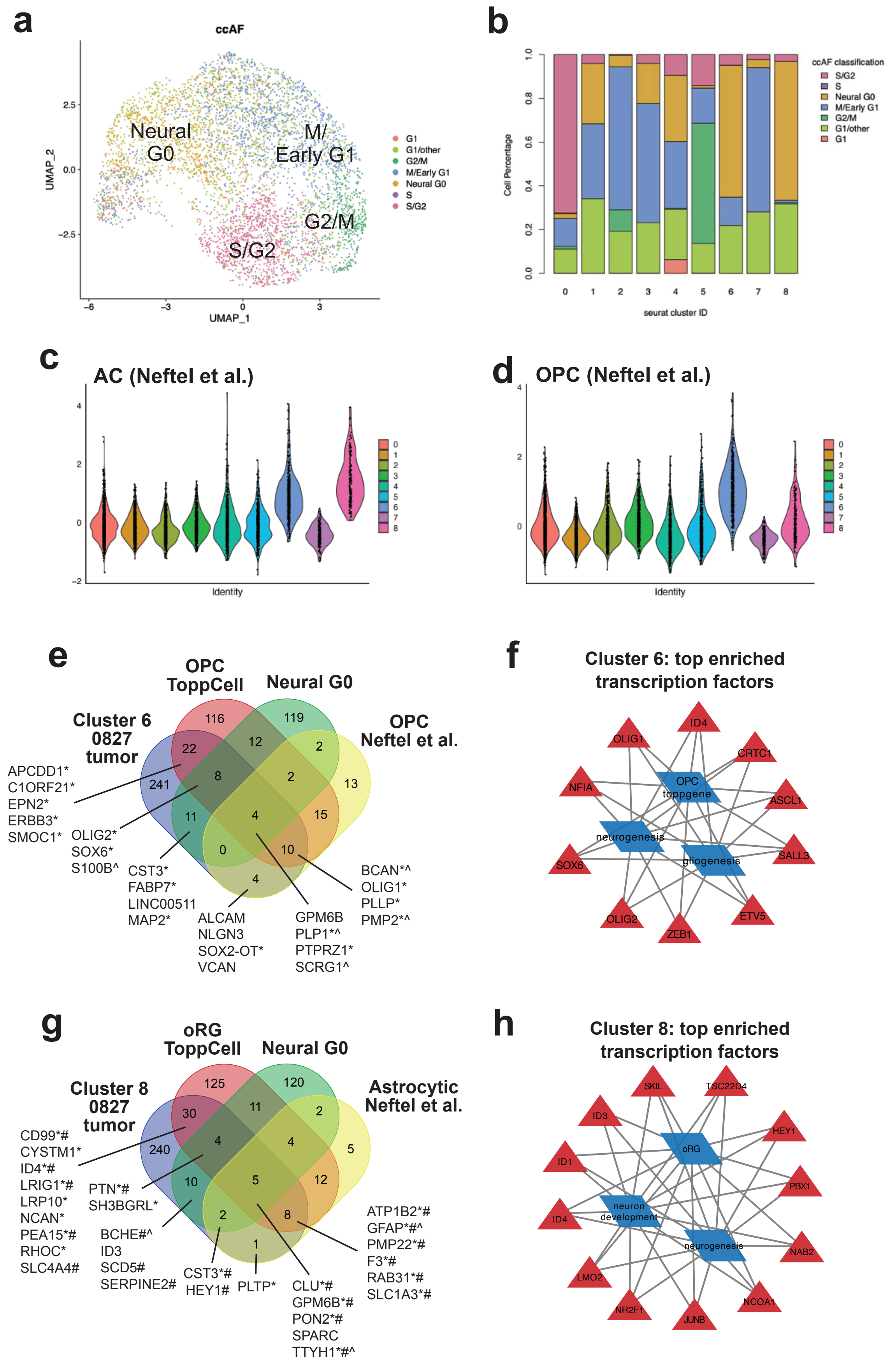

**Extended Data Figure 3**: Gene expression analysis of cells of clusters for the GSC-0827 tumor reference associated with **Figure 1**.
**a**, Cell cycle phase classification of each cell in GSC-0827 tumor reference using ccAF (O'Connor et al., 2021). The Neural G0 state designation is derived from a human NSC G0-like state that contains a mixture of genes expressed in adult quiescent NSCs, fetal radial glial (RG) cells, and oligodendrocyte progenitor (OPC) cells (O'Connor et al., 2021).
**b**, Analysis from a showing percent make up of each cluster. Clusters 6 and 8 show the highest percentages of Neural G0 cells.
**c-d**, Gene expression module scores for clusters in in GSC-0827 tumor reference using GBM gene expression modules derived from Neftel et al., 2019.
**e-h**, Analysis of gene enriched in clusters 6 and 8 of GSC-0827 tumor reference associated with Figure 1.
**e**, Overlap of top 300 enriched genes from UMAP cluster 6 (OPC) of GSC-0827 tumor reference (Figure 1) with ToppCell Non-dividing OPC gene cluster genes, Neural G0 genes, and the OPC GBM genes from Neftel et al, 2019. *OPC expression from Chamling et al., 2021 (PMID: 33510160). ^Indicates presence among 22 common GBM-specific Neural G0 marker genes (O'Connor et al., 2021).
**f**, A network of Cluster 6 enriched transcription factors showing associations with Toppgene Non-dividing
OPC gene cluster, and gene ontology terms neurogenesis (GO:0022008) and gliogenesis (GO:0042063).
**g**, Overlap of top 300 enriched genes from UMAP cluster 6 (oRG/Ac) of GSC-0827 tumor reference (Figure 7A) with oRG ToppCell cluster genes, Neural G0 genes, and the Astrocytic GBM genes from Neftel et al, 2019.
*Indicates expression >2-fold enriched in oRG cells from Nowakowski et al., 2017 (PMID: 29217575). #Indicates expression >2-fold enriched in astrocytes expression from Nowakowski et al., 2017 (PMID: 29217575).
**h**, A network of Cluster 8 enriched transcription factors showing associations with ToppCell Radial_glial-oRG gene cluster, and gene ontology terms neuron development (GO:0048666) and neurogenesis (GO:0022008). ^Indicates presence among 22 common GBM-specific Neural G0 marker genes (O'Connor et al., 2021).

A full gene list for each Venn diagram is available in Supplementary Table 10.

### Extended Data Figure 4

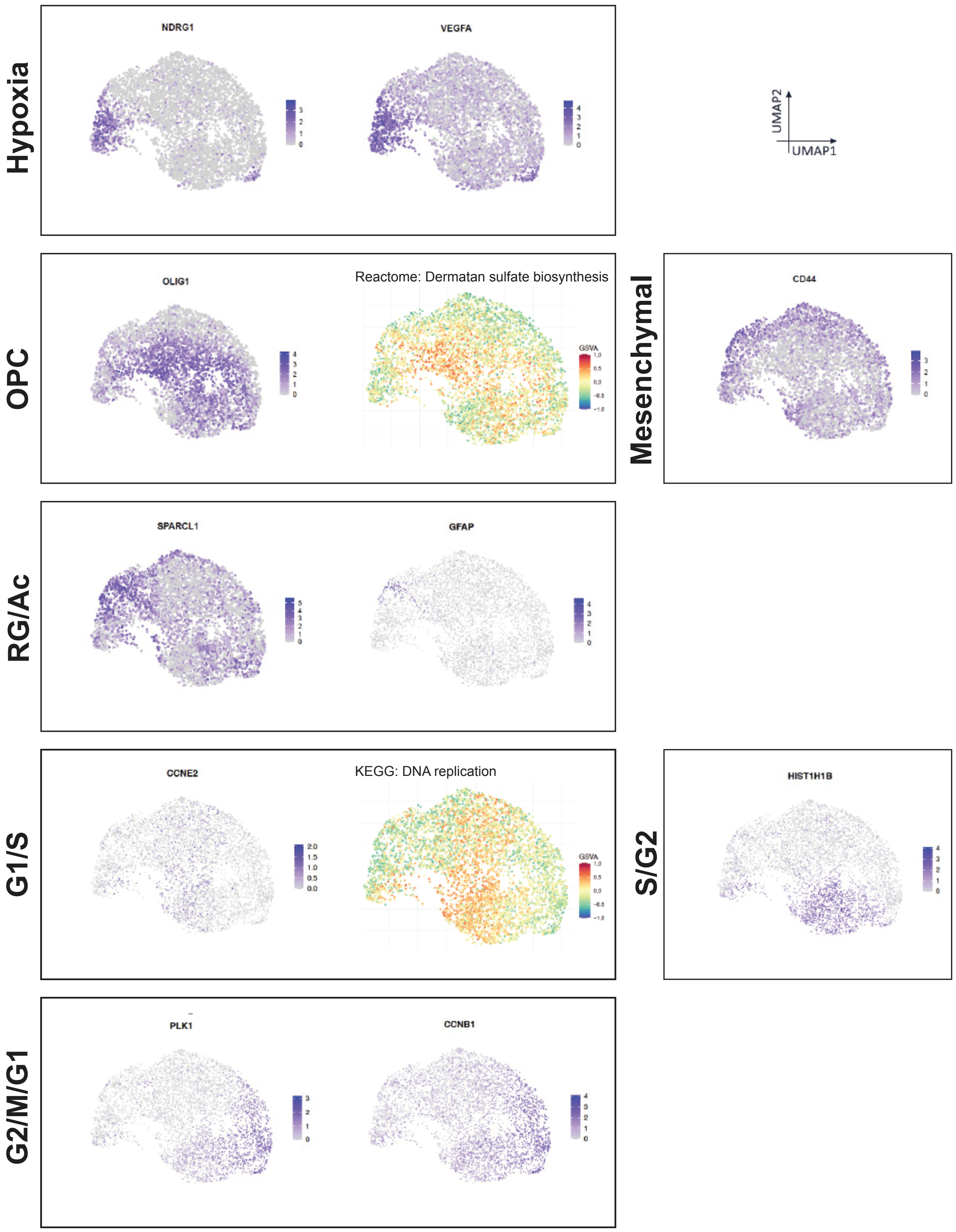

**Extended Data Figure 4**: Gene expression analysis of cells of clusters for the GSC-0827 tumor reference associated with **Figure 1**. Gene expression analysis of individual genes and Gene Set Variation Analysis (GSVA) associated with clusters and cell cycle phases for GSC-0827 tumor reference associated with Figure 1. Note that Reactome: Dermatan sulfate biosynthesis includes category includes BCAN, CSPG4, CSPG5, NCAN, and VCAN, which are all enriched in cluster 6 (OPC).

### Extended Data Figure 5

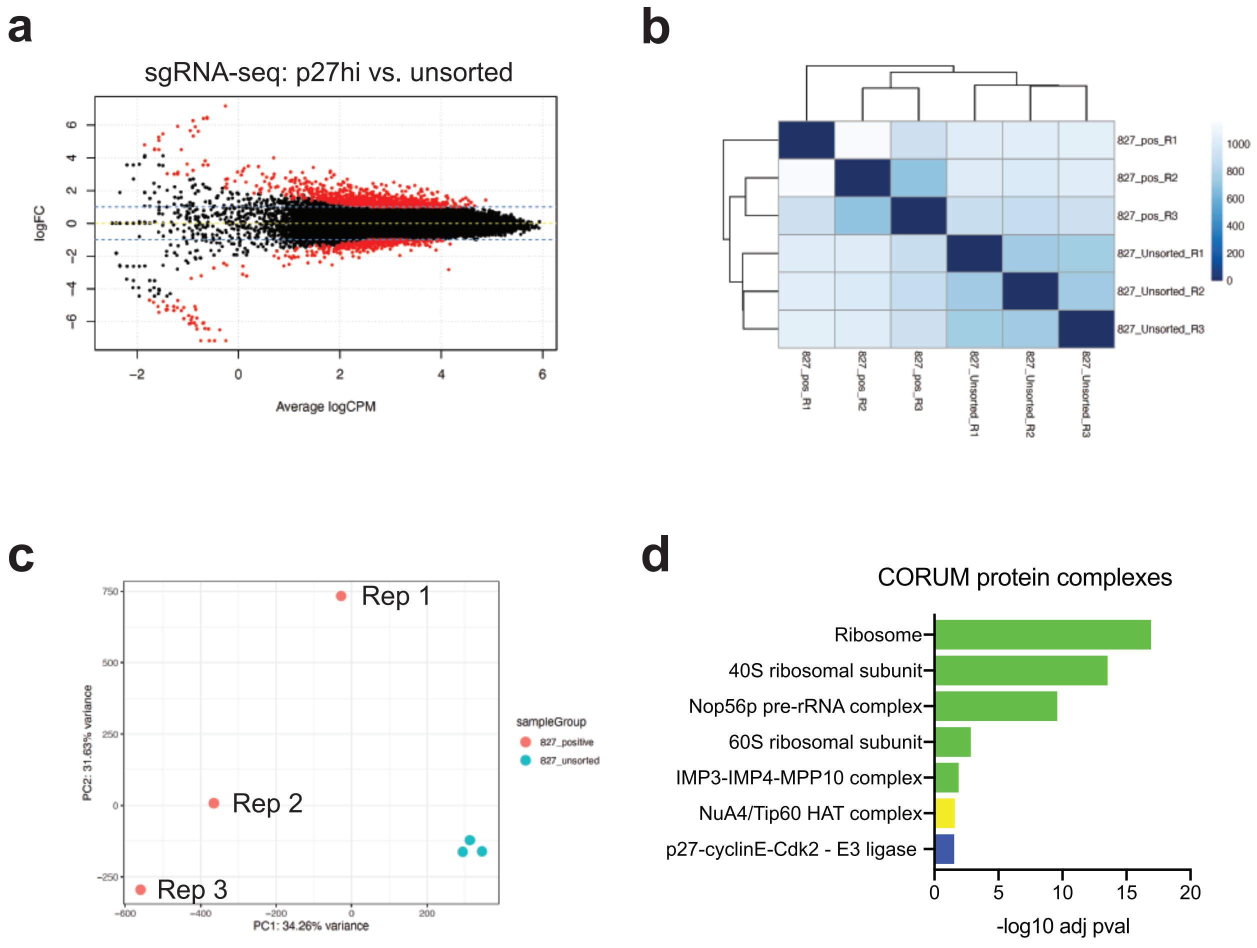

**Extended Data Figure 5**: Data supporting **Figure 3**.
**a**, M (log fold change) versus A (mean expression) plot from sgRNA-seq for G0-trap screen. Red indicates positively scoring enriched or depleted sgRNA.
**b**, Correlation plot of screen replicates.
**c**, Principal component analysis of screen replicates.
**d**, Gene set enrichment analysis of screen hits using CORUM protein complex database (http://mips.helmholtz-muenchen.de/corum/).

### Extended Data Figure 6

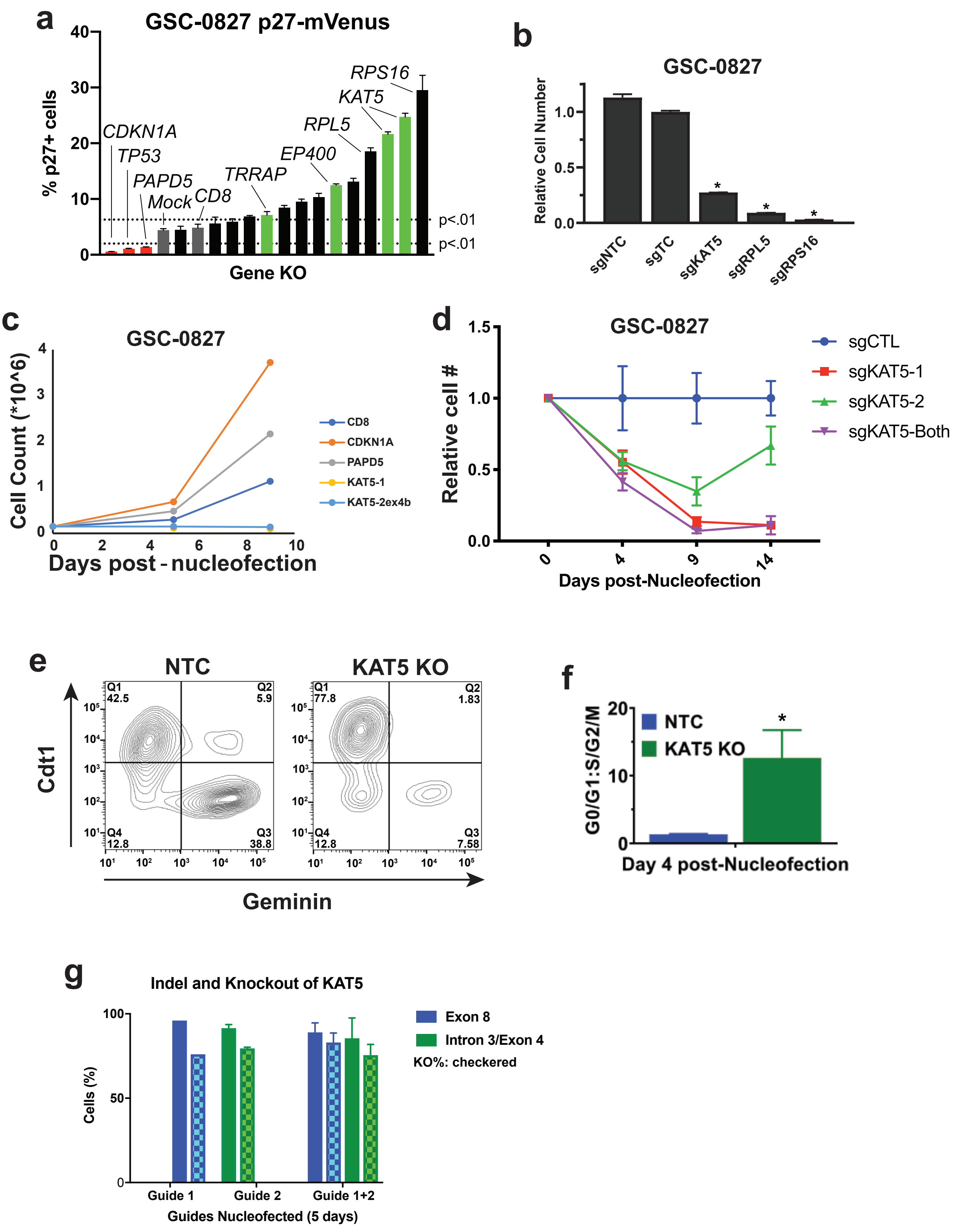

**Extended Data Figure 6**: KAT5 inhibition triggers G0-like state in GSCs in *vitro*, in support of **Figure 3**.

**a**, Select retest of G0-trap screen hits using nucleofection of sgRNA:Cas9 RNPs in GSC-0827 p27-mVenus cells. Cells were flow analyzed 5 days post-nucleofection (n=3; student's t-test, p<.01). This technique allows for penetrant KO of target genes in GSCs in a relatively short period time (~48hrs) (Hoellerbauer et al., 2020a).

**b**, Comparison cell number in p53 KO and control GSC-0827 cells after KAT5, RPL5, and RPS16 inhibition (n=3; student's t-test, p<.01).

**c**, Cell counts at various times post-nucleofection of KAT5 and two negative screen hits found significantly depleted in GSC-0827 $p27^{hi}$ cells, CDKN1A and PAPD5.

**d**, Outgrowth of GSC-0827 cells after nucleofection of sgKAT5:Cas9 or sgCTRL:cas9 RNP complexes (n=3; student's t-test, p<.01).

**e**, Flow analysis of FUCCI factors in GSC-0827 cells 5 days post-nucleofection of sgKAT5:Cas9 or sgCTRL:Cas9 RNP complexes.

**f**, Quantification of G0/G1:G2/M ratios from **e** (n=3; student's t-test, p<.01).

**g**, Analysis of indel efficiency and knockout of KAT5 using sgRNA:Cas9 RNPs in GSC-0827 cells.

### Extended Data Figure 7

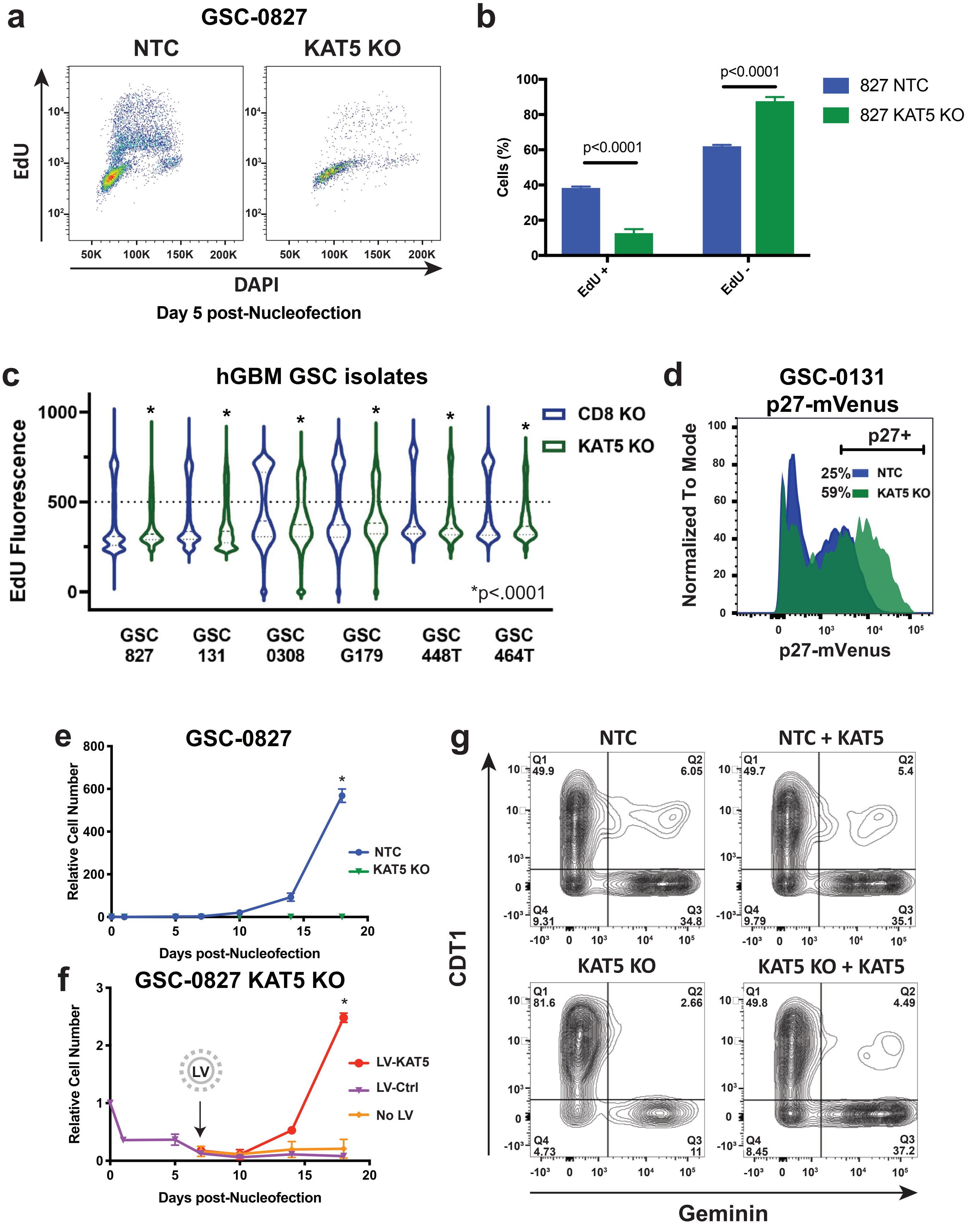

**Extended Data Figure 7**: KAT5 inhibition triggers G0-like state in GSCs in *vitro*, in support of **Figure 3**.
**a**, FACS-based assessment of EdU incorporation after KAT5 KO using nucleofection of sgRNA:Cas9 RNPs (5 days post-nucleofection) in GSC-0827 cells.
**b**, Quantification of loss of EdU incorporation for this experiment (n=3; student's t-test, p<.01).
**c**, Violin plots of 3hrs of EdU incorporation KAT5 KO 5 days post-nucleofection in various GSC isolates. KS test was used to test significance (p<.0001).
**d**, Retest of KAT5 inhibition in GSC-0131 p27-mVenus cells using nucleofection of sgRNA:Cas9 RNPs. Cells were flow analyzed 5 days post-nucleofection.
**e**, Evaluation of growth of KAT5 KO GSC-0827 cells at days shown post-nucleofection. (n=3; student's t-test, p<.001).
**f**, KAT5 KO rescue and reversibility experiment. To examine reversibility of KAT5 KO growth defect, we first knocked out KAT5 using sgKAT5:Cas9 in GSC-0827 cells and then added KAT5 back 7 days later via lentiviral transduction as an expressed KAT5 ORF. By 7 days, the cells that had received the LV-KAT5 ORF had growth rates return to near parental cell levels while control LV cells were no different than KAT5 KO in **e**.
**g**, Flow analysis of FUCCI factors after rescue 14 days post-nucleofection and 10 days post-rescue. After 10 days, the LV-KAT5 ORF cells had their cell cycle phase ratios return to parental cell levels, as judged by FUCCI reporter analysis, while the LV-control cells remained similar to KAT5 KO.

### Extended Data Figure 8

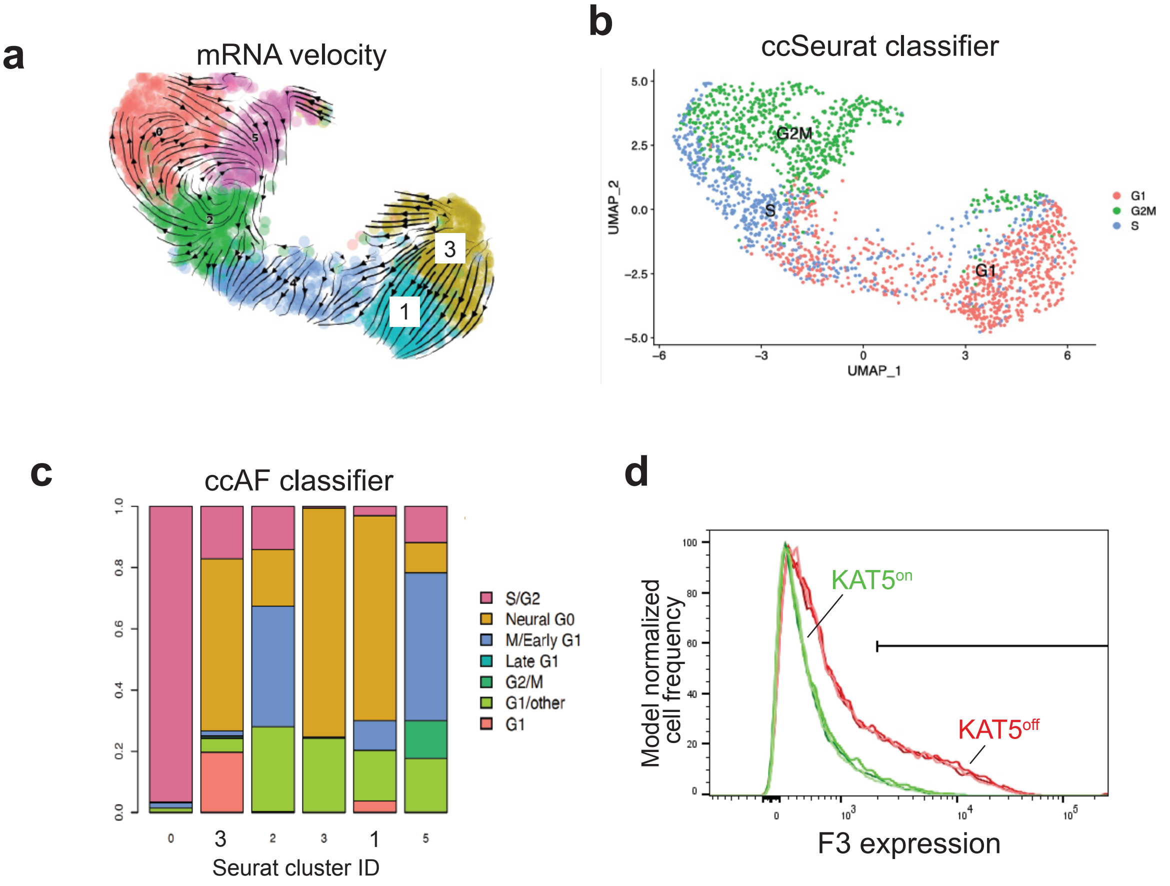

**Extended Data Figure 8**: Single cell gene expression analysis of KAT5 KO in GSC-0827 cells in vitro, in support of **Figure 3**.
**a**, RNA velocity analysis of scRNA-seq data from **Figures 3e-f**.
**b**, Cell cycle state predictions using the ccSeurat classifier for scRNA-seq data from **a**.
**c**, Cell cycle phase predictions using the ccAF classifier for scRNA-seq data from **a**.
**d**, F3 expression evaluated by flow analysis of GSC-0827 C13 cells grown in the presence of Dox ($KAT5^{on}$) or 5 days post withdrawal ($KAT5^{off}$).

### Extended Data Figure 9

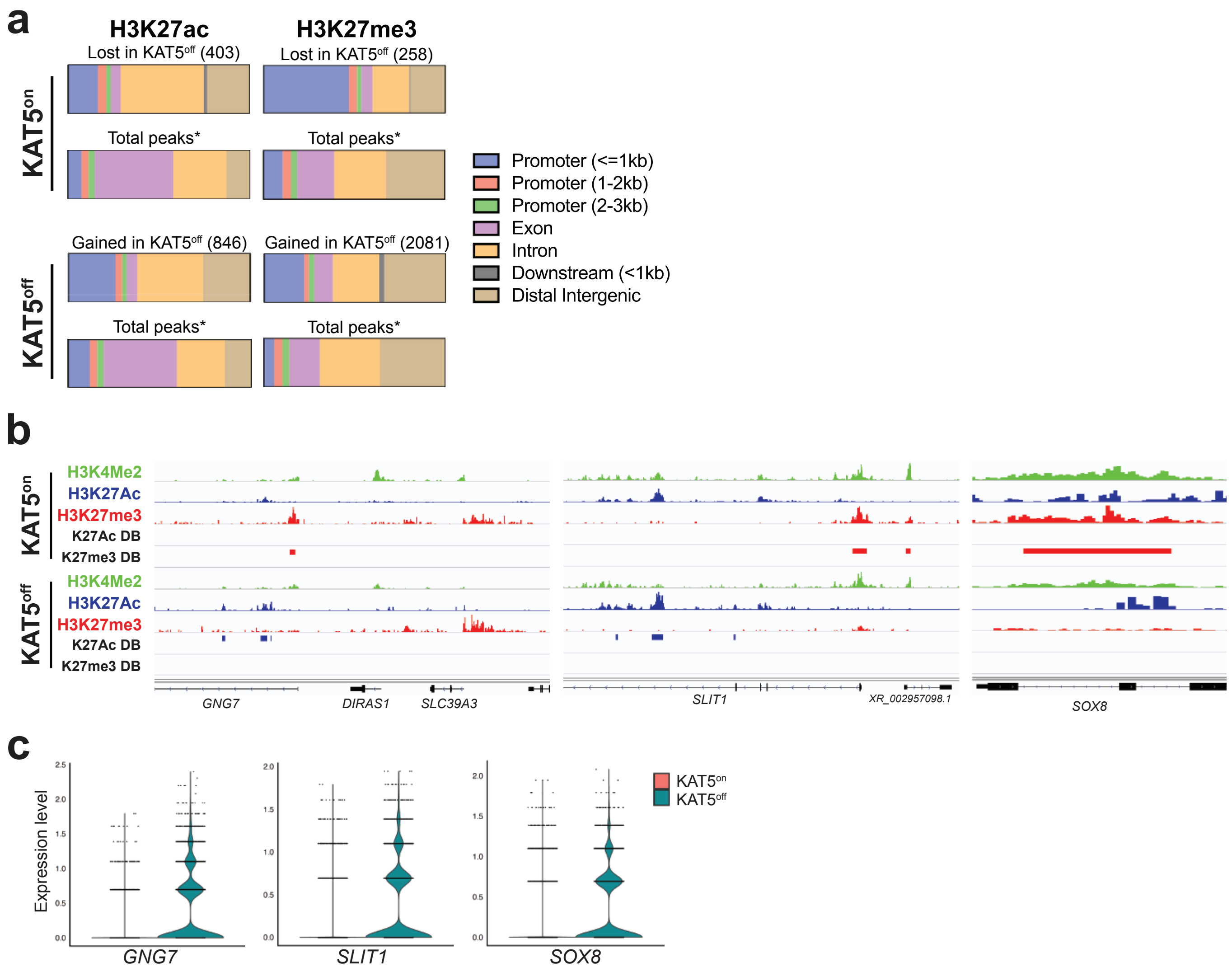


## d

KAT5$^{on}$ Superenhancers

| Chrom. | Start | End | Width | Slope | Gene | Description | Relevance |
|---|---|---|---|---|---|---|---|
| chr7 | 13032102 | 13063402 | 31301 | 1563.6 | ***ARL4A*** | ADP ribosylation factor like GTPase 4A | glioma-associated antigen |
| chr7 | 54813258 | 54813492 | 235 | 627.5 | ***SEC61G*** | SEC61 translocon subunit gamma | GBM: survival / ER stress |
| chr7 | 101154436 | 101167265 | 12830 | 489.1 | ***VGF*** | VGF nerve growth factor inducible | GBM: stemness / survival |
| chr6 | 150753222 | 150775852 | 22631 | 189.8 | ***MTHFD1L*** | methylenetetrahydrofolate dehydrogenase 1 like | folate enzyme / oncogenesis |
| chr1 | 164576043 | 164612598 | 36556 | 82.4 | ***PBX1*** | PBX homeobox 1 | SVZ stem and progenitor cells |
| chr19 | 44742168 | 44753542 | 11375 | 72.0 | ***BCL3*** | BCL3 transcription coactivator | GBM: EMT/ NF-kB |
| chr1 | 183000000 | 183000000 | 50644 | 48.3 | ***RGS16*** | regulator of G protein signaling 16 | GBM: mesenchymal |
| chr7 | 92755155 | 92769276 | 14122 | 48.2 | ***CDK6*** | cyclin dependent kinase 6 | GBM: G1/S transition |
| chr11 | 35151288 | 35166564 | 15277 | 25.5 | ***CD44*** | CD44 cell-surface glycoprotein | GBM: mesenchymal |

## e

KAT5$^{off}$ Superenhancers

| Chrom. | Start | End | Width | Slope | Gene | Description | Relevance |
|---|---|---|---|---|---|---|---|
| chr1 | 230728655 | 230747162 | 18508 | 18507.0 | ***AGT*** | angiotensinogen | GBM: bevacizumab biomarker |
| chr1 | 156406287 | 156421983 | 15697 | 15696.0 | ***C1ORF61*** | MIR9-1 host gene | Brain development / oncogenesis |
| chr17 | 44907324 | 44917306 | 9983 | 9982.0 | ***GFAP*** | glial fibrillary acidic protein | Astrocytes and astryocytoma |
| chr19 | 2685571 | 2696991 | 11421 | 552.3 | ***GNG7*** | G protein subunit gamma 7 | GBM: expression |
| chr3 | 47536092 | 47554182 | 18091 | 104.8 | ***CSPG5*** | chondroitin sulfate proteoglycan 5 | OPC progenitors |
| chr2 | 119370840 | 119387743 | 16904 | 23.1 | ***DBI*** | diazepam binding inhibitor, acyl-CoA binding protein | quiescent cancer cells |
| chr1 | 156638190 | 156655316 | 17127 | 15.9 | ***BCAN*** | brevican | GBM: expression / invasiveness |
| chr14 | 69963927 | 69964272 | 346 | 9.2 | ***SMOC1*** | SPARC-related modular calcium-binding protein 1 | Glioma/GBM/OPC: expression |
| chr16 | 986231 | 991762 | 5532 | 2.1 | ***SOX8*** | SRY-box transcription factor 8 | OLIG1+ low grade glioma |

**Extended Data Figure 9**: Gene all outs of genomic regions from CUT&TAG analysis of GSC-0827 Clone 13 tumors, in support of **Figure 5**.
**a**, Comparison of distribution of H3K27ac and H3K27me3 peaks from tumors among genomic regions for peaks significantly different in KAT5$^{on}$ and KAT5$^{off}$ conditions. Total significant peaks shown in parentheses; *MACS2 called peaks >11,000 for ac and >47,000 for me3.
**b**, Additional examples of genes with altered significantly altered H3K27ac/me3 peaks in KAT5$^{on}$ and KAT5$^{off}$ C13 tumors.
**c**, Gene expression analysis from scRNA-seq data for KAT5$^{on}$ and KAT5$^{off}$ C13 tumors.
**d**, Call outs of super enhancers among 521 predicted from KAT5$^{on}$ GSC-0827 tumors.
**e**, Call outs of super enhancers among 218 predicted from KAT5$^{off}$ GSC-0827 tumors.
Complete data sets can be found in **Supplementary Table 15**.

### Extended Data Figure 10

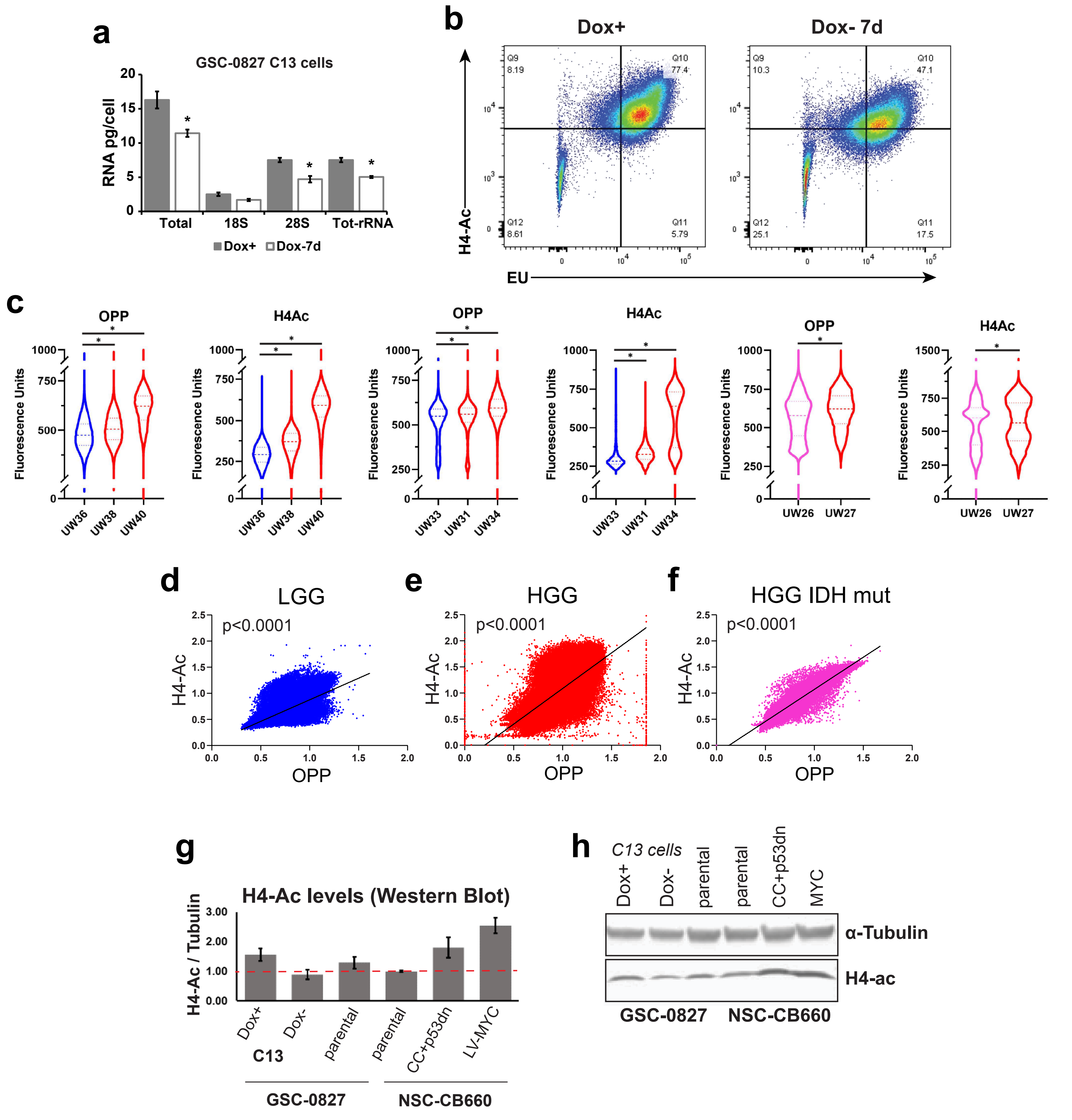

**Extended Data Figure 10**: Data in support of **Figure 7**, including analysis of RNA levels and EU incorporation and assessment of KAT5 activity and protein translation rates in primary glioma tumor samples.

**a**, Total RNA and rRNA quantification using TapeStation RNA assay system (Agilent) using RNA isolated from GSC-0827 C13 cells grown in Dox+ and Dox- conditions (7 days).

**b**, FACS analysis of histone H4 acetylation and p27 levels after 0 and 7 days of Dox withdrawal in C13 cells. EU treated was performed for 30mins before harvest.

**c**, Violin plots of OPP and pan-H4-Ac assay results for 2 LGG (IDH1/2mut) (blue), 5 HGG (IDH1/2wt) (red), and 1 HGG (IDH1/2 mut) (pink) tumors. KS test was used to assess significance ($p<0.0001$)

**d-f**, Combined of OPP versus pan-H4-Ac for LGG (Pearson r = 0.77), HGG (Pearson r =0.68), and HGG (IDH1/2mut) tumors (Pearson r = 0.93), respectively; $p<0.0001$ for each.

**g**, Quantification of Western blot results shown in **h** for H4-Ac (pan) for GSC-0827 and NSC-CB660 cells. C13 = clone 13 Dox-KAT5 cells (Dox+ indicates continuous Dox+ growth; Dox- indicates doxycycline removal for 4 days). CC+p53dn= CyclinD1+ CDK4R24C+ dominant-negative p53+TERT (Hubert et al., 2013).

### Supplementary Figure 1

**a**

**Tumor reference**

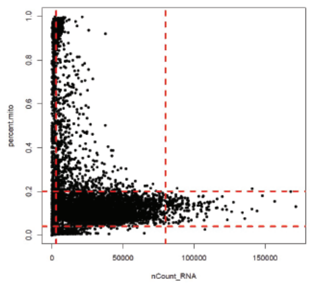


Before filtering = 8404
Tumor 1 = 1458 cells
Tumor 2 = 2016 cells
Tumor 3 = 4930 cells
Filters applied :
0.2 < percent.mito < 0.04
3000 < nCount_RNA < 80000
After filtering = 5144

**b**

**p27$^{hi}$ tumor cells**

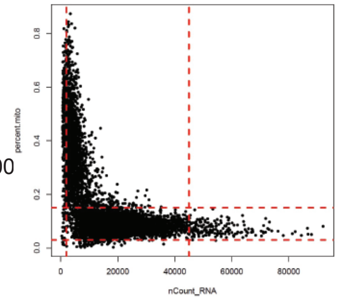


Before filtering = 7338 cells
Filters applied:
0.03 < percent.mito < 0.15
2000 < nCount_RNA < 45000
After filtering = 4780 cells

**c** **d** **e** **f**

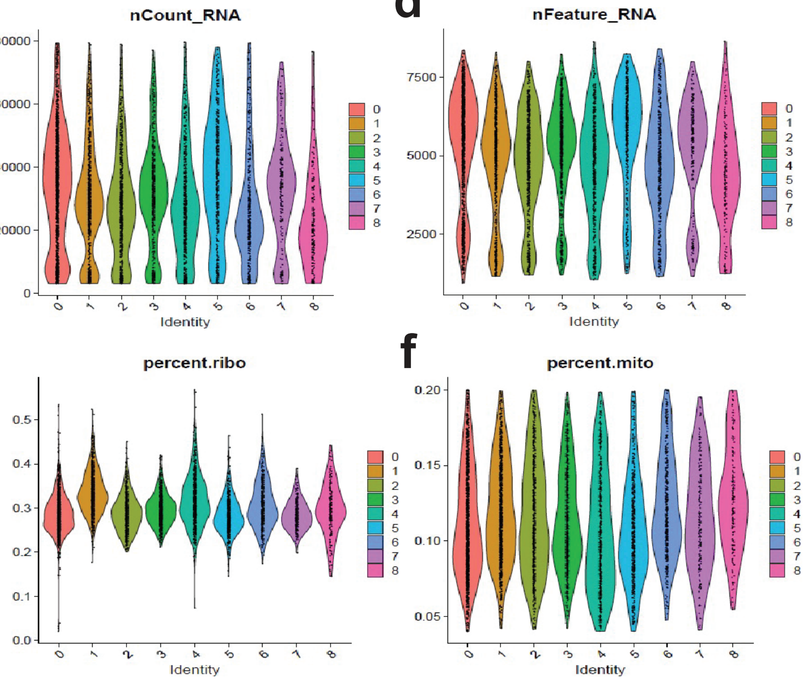

**Supplementary Figure 1**: scRNA-seq filtering and data quality evaluation for **Figure 1**.
**a**, Experimental overview for collection of samples for scRNA-seq.
**b**, Filtering criteria used for each scRNA-seq data set.
**c-f**, scRNA-seq quality control metrics viewed by cluster after filtering of data.

### Supplementary Figure 2

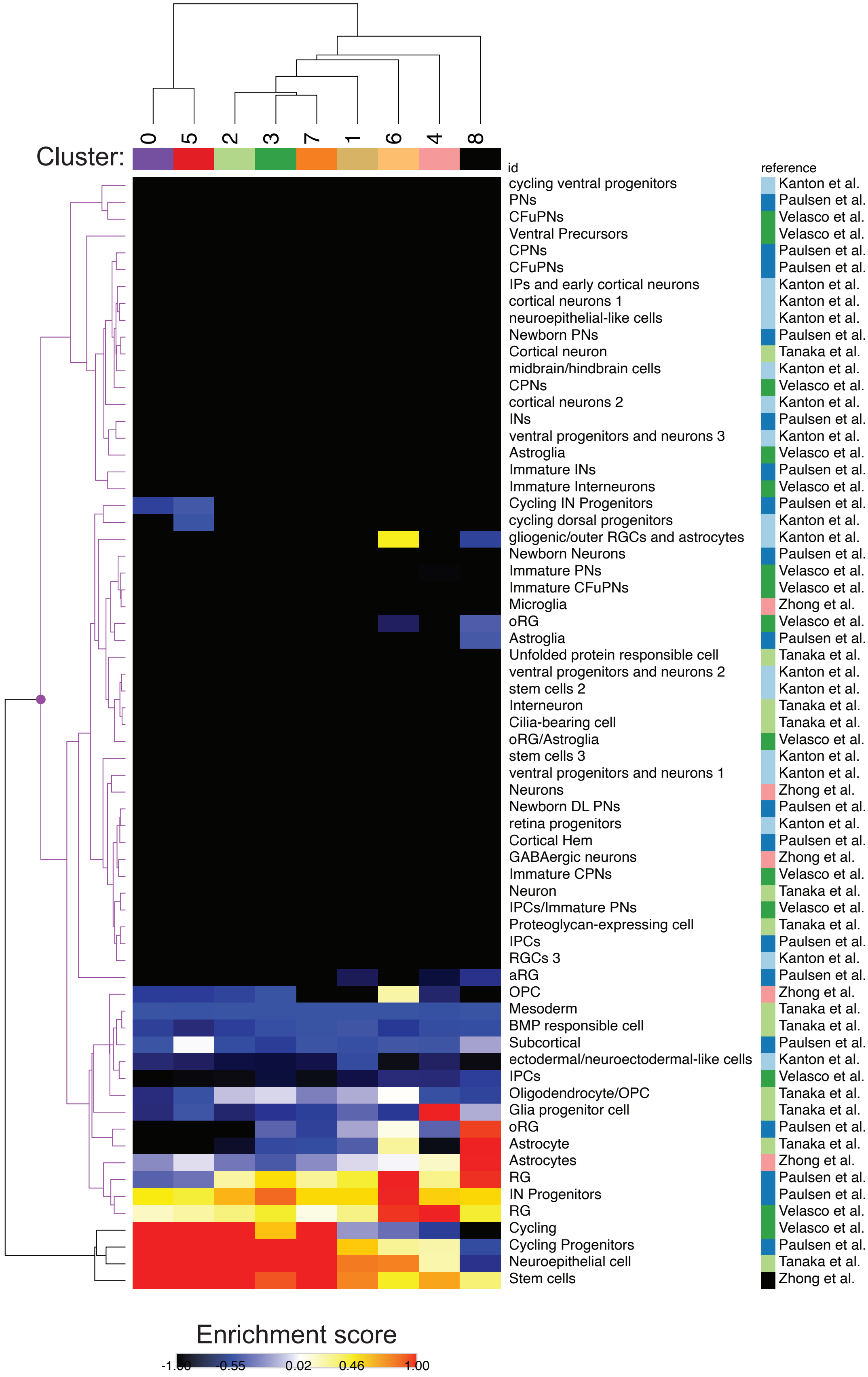

**Supplementary Figure 2**: The full heatmap of neurodevelopmental gene expression profiles from **Figure 1h** for human fetal and adult neurodevelopmental cell types in the scRNA-seq ToppCell Atlas. Tumor cluster gene sets to enriched in tumor clusters to those for neurodevelopmental cell types available in the ToppCell Atlas[1] using linear regression analysis[2]. ToppCell Atlas data and cell descriptions are taken from four references shown on the right-side column[3-7].


1 Jin, K. *et al.* An Interactive Single Cell Web Portal Identifies Gene and Cell Networks in COVID-19 Host Responses. *iScience*, 103115, doi:10.1016/j.isci.2021.103115 (2021).

2 Young, M. D. *et al.* Single-cell transcriptomes from human kidneys reveal the cellular identity of renal tumors. *Science* **361**, 594-599, doi:10.1126/science.aat1699 (2018).

3 Kanton, S. *et al.* Organoid single-cell genomic atlas uncovers human-specific features of brain development. *Nature* **574**, 418-422, doi:10.1038/s41586-019-1654-9 (2019).

4 Tanaka, Y., Cakir, B., Xiang, Y., Sullivan, G. J. & Park, I. H. Synthetic Analyses of Single-Cell Transcriptomes from Multiple Brain Organoids and Fetal Brain. *Cell Rep* **30**, 1682-1689 e1683, doi:10.1016/j.celrep.2020.01.038 (2020).

5 Velasco, S. *et al.* Individual brain organoids reproducibly form cell diversity of the human cerebral cortex. *Nature* **570**, 523-527, doi:10.1038/s41586-019-1289-x (2019).

6 Zhong, S. *et al.* A single-cell RNA-seq survey of the developmental landscape of the human prefrontal cortex. *Nature* **555**, 524-528, doi:10.1038/nature25980 (2018).

7 Paulsen, B. *et al.* Autism genes converge on asynchronous development of shared neuron classes. *Nature* **602**, 268-273, doi:10.1038/s41586-021-04358-6 (2022).

### Supplementary Figure 3

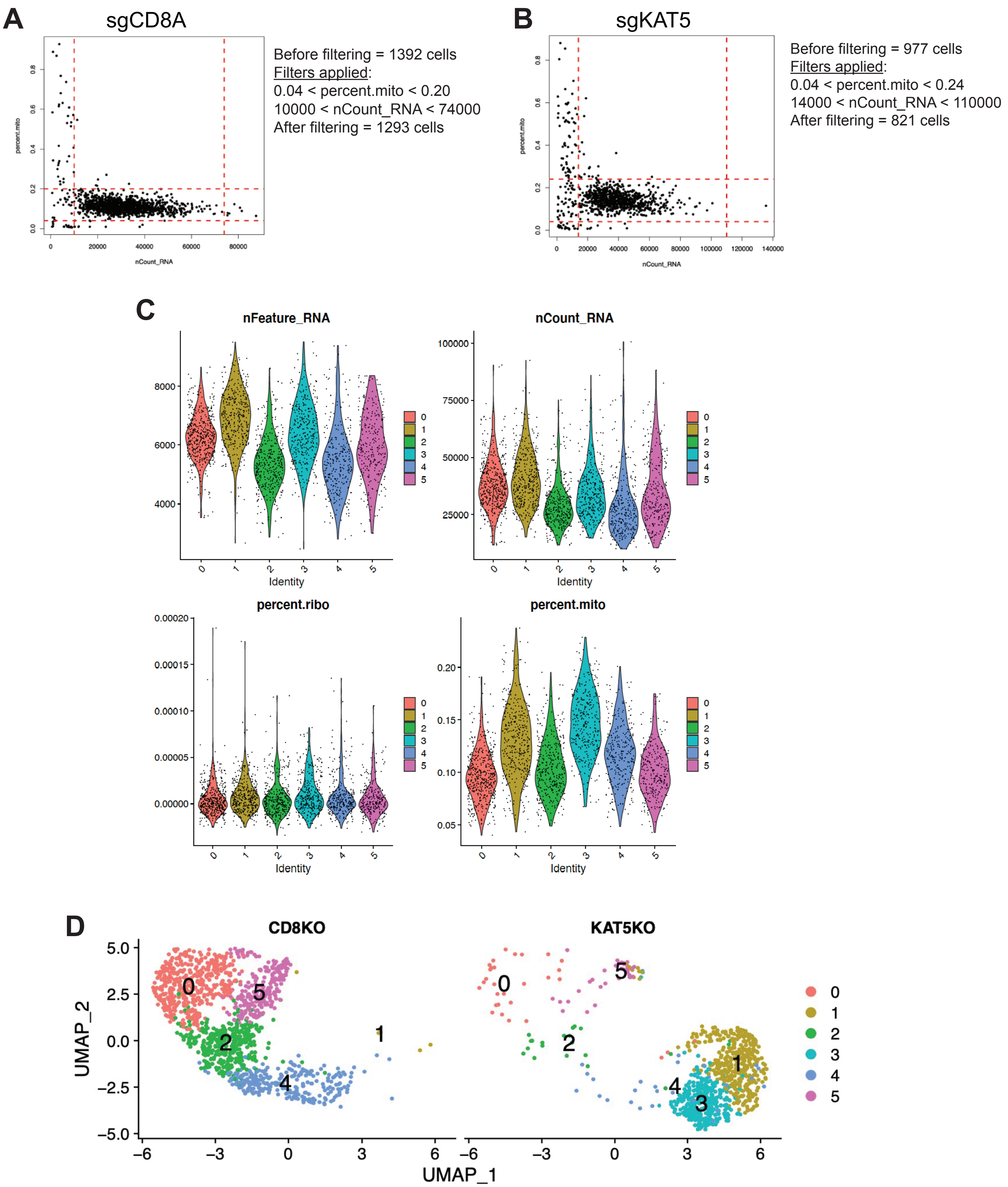

**Supplementary Figure 3**: scRNA-seq filtering and data quality evaluation for **Figure 3**.
**a-b**, Filtering criteria used for sgCD8A and sgKAT5 GSC-0827 scRNA-seq analysis.
**c**, Visualization of scRNA-seq quality control parameters for co-embedded KAT5 and CD8 KO GSC-0827 scRNA-seq data showing cluster scheme from **Figure 3**.
**d**, Individual UMAP projections of the co-embedded for co-embedded KAT5 and CD8 KO GSC-0827 scRNA-seq data from **Figure 3**.
